# Unveiling the influence of tumor and immune signatures on immune checkpoint therapy in advanced lung cancer

**DOI:** 10.1101/2024.04.15.589544

**Authors:** Nayoung Kim, Sehhoon Park, Areum Jo, Hye Hyeon Eum, Hong Kwan Kim, Kyungjong Lee, Jong Ho Cho, Bo Mi Ku, Hyun Ae Jung, Jong-Mu Sun, Se-Hoon Lee, Jin Seok Ahn, Jung-Il Lee, Jung Won Choi, Dasom Jeong, Minsu Na, Huiram Kang, Jeong Yeon Kim, Jung Kyoon Choi, Hae-Ock Lee, Myung-Ju Ahn

## Abstract

This study investigates the variability among patients with non-small cell lung cancer (NSCLC) in their responses to immune checkpoint inhibitors (ICI). Recognizing that patients with advanced-stage NSCLC rarely qualify for surgical interventions, it becomes crucial to identify biomarkers that influence responses to ICI therapy. We conducted an analysis of single-cell transcriptomes from 33 lung cancer biopsy samples, with a particular focus on 14 core samples taken before the initiation of palliative ICI treatment. Our objective was to link tumor and immune cell profiles with patient responses to ICI. We discovered that ICI non-responders exhibited a higher presence of CD4+ regulatory T cells, resident memory T cells, and TH17 cells. This contrasts with the diverse activated CD8+ T cells found in responders. Furthermore, tumor cells in non-responders frequently showed heightened transcriptional activity in the NF-kB and STAT3 pathways, suggesting a potential inherent resistance to ICI therapy. Through the integration of immune cell profiles and tumor molecular signatures, we achieved an discriminative power (AUC) exceeding 95% in identifying patient responses to ICI treatment. These results underscore the crucial importance of the interplay between tumor and immune microenvironment, including within metastatic sites, in affecting the effectiveness of ICIs in NSCLC.

## Introduction

Treatment landscape in cancer has rapidly evolved with the introduction of immune checkpoint inhibitors (ICI). Among various immune checkpoints, antibody-targeting programmed cell death-1 (PD-1) and its ligand (PD-L1) have demonstrated clinical benefits over conventional systemic chemotherapy in patients with non-small cell lung cancer (NSCLC). They have been approved as either monotherapy in patients with high PD-L1 expression ^1^ or combined with cytotoxic chemotherapy regardless of PD-L1 expression ^2,3^. Moreover, the clinical benefits were validated in unresectable stage III NSCLC as consolidation therapy after definitive chemoradiotherapy or early stage NSCLC (Ib-IIIA) as adjuvant therapy after curative surgery ^4^.

There have been many efforts to elucidate the predictive biomarkers of PD-(L)1 inhibitors. The PD-L1 expression in tumor tissue evaluated via immunohistochemistry has been incorporated as a companion diagnostic biomarker from the early clinical trials, which enhanced the response rate up to 46% in patients with PD-L1 ≥50% and showed an overall survival rate of up to 26.3 months ^5^. Other biomarkers, such as tumor mutation burden or gene expression profile, also demonstrated a positive predictive value ^6,7^. Recent large-scale meta-analysis of clinico-immunogenomics shows that both tumor- and T cell intrinsic factors exert a substantial impact on ICI response ^8^, supporting the necessity of in-depth investigation of high-throughput profiles of tumor and microenvironment.

ICI treatment modifies systemic immune responses, which can be monitored by alterations in the proportion of specific immune cell populations. An increase in PD-1+Ki67+CD8+ T cells in peripheral blood after PD-1 inhibitor treatment is associated with better outcomes in patients with NSCLC ^9,10^. This cell type has also been identified in tumor tissues prior to the ICI treatment, where it is linked to clinical outcomes, and shows impaired production of classical effector cytokines ^11^. More specifically, PD-1 expression is upregulated upon antigen recognition ^12^, indicating that certain T cells in the tumor microenvironment are actively engaged as tumor-specific T cells. Beyond CD8+ T cells, other immune cell types, such as myeloid-derived suppressor cells or regulatory T cells, which regulate tumor-specific T cell immunity may also influence therapeutic outcomes ^13–15^. Overall, the multicellular regulation of the tumor-immune microenvironment emphasizes the importance of profiling systemic tumor and immune cells at single cell resolution to investigate factors associated with responses to ICI treatment.

## Results

### Variability and features of the lung cancer samples

We conducted scRNA-seq on 33 lung cancer samples from 26 patients treated with immune checkpoint inhibitors (ICI) between August 2017 and December 2019 to understand how cellular dynamics in lung cancer affect treatment sensitivity to PD-(L)1 inhibitors, used alone or in combination (Fig. 1a and Table S1). Immune checkpoint therapy provides clinical benefit in advanced metastatic NSCLC across different treatment lines ^16^. Notably, our samples have been collected from various tissue sites. In the scRNA-seq analysis, all specimens were used for the cell type profiling in an unbiased manner. For the evaluation of clinical outcomes, only refined 14 core samples from 11 patients were used to minimize sample specific variations. Exclusion criteria from the core group encompass samples with treatment applied as adjuvant therapy, acquired after ICI treatment, no tumor content, non-evaluable for the clinical response, or histology other than non-small cell lung cancer such as nuclear protein in testis (NUT) and small cell lung cancer (SCLC) (Table 1 and Fig. 1a). Of the 11 core patients, 8 had adenocarcinoma (ADC) and 3 had squamous cell carcinoma (SQ). Clinical outcomes of ICI were partial response (PR) in four patients, stable disease (SD) in two patients, and progressive disease (PD) in five patients. Patients were classified as responders (PR) and non-responders (SD and PD) according to ICI response.

**Figure 1.**
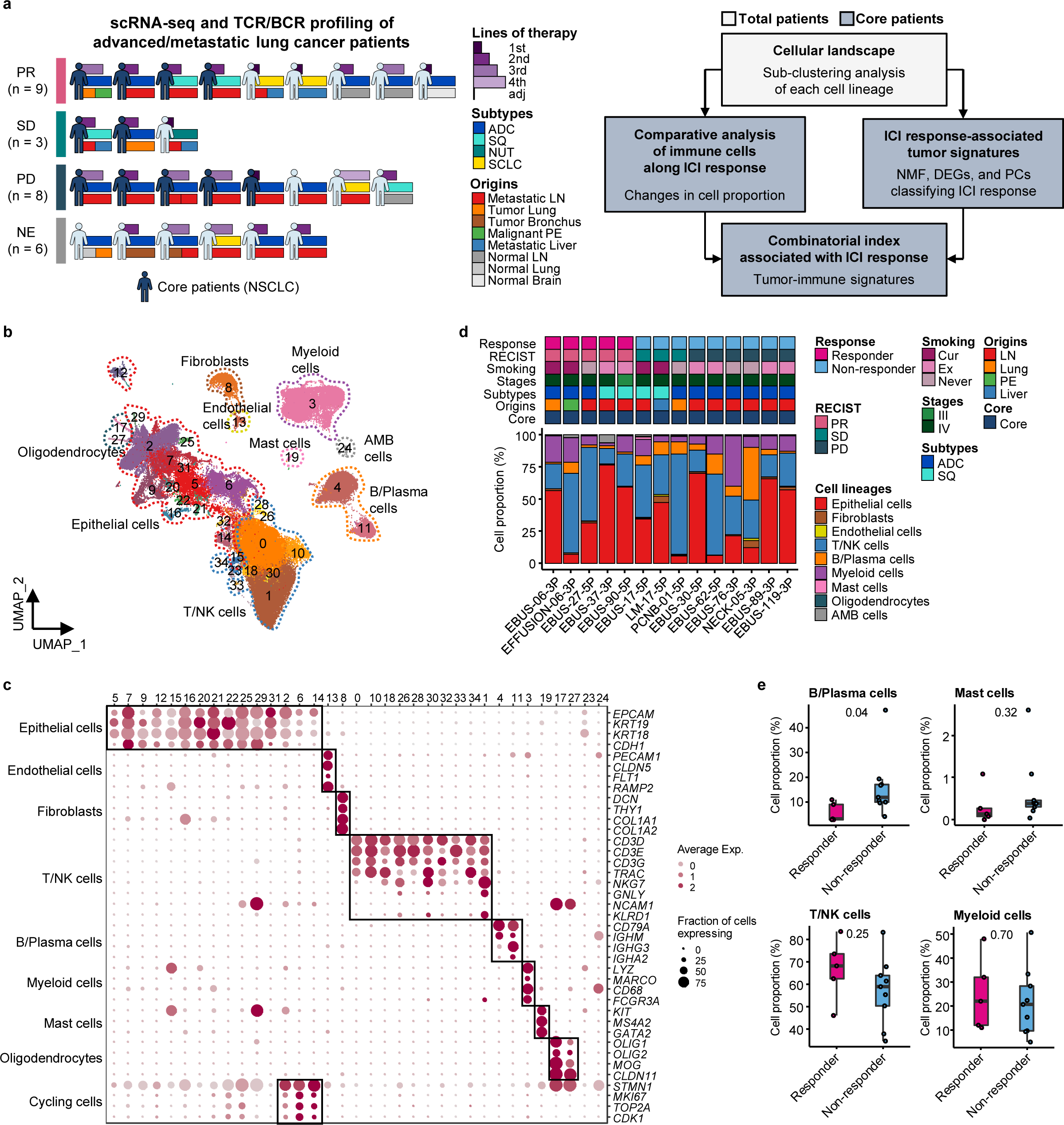
Cell lineage identification of 96,505 single cells from 26 patients with lung cancer treated with ICI. **a,** Workflow of sample collection and single-cell analysis of lung cancer patients treated with ICI. **b,** UMAP plot of 96,505 single cells from 33 samples acquired from 26 advanced lung cancer patients, colored by clusters. AMB cells, Ambiguous cells. **c,** Dot plot of mean expression of canonical marker genes for cell lineages. **d,** Proportions of the cell lineages in NSCLC tissue from core patients shown by individual samples aligned with clinical data. Labels for origins indicate LN, Metastatic lymph node; Lung, Tumor lung; PE, Malignant pleural effusion; Liver, Metastatic liver. **e,** Box plot of the percentage of cell lineages in responder and non-responder groups. Label represents p-value calculated via two-tailed Student’s t-test. Each box represents the median and the interquartile range (IQR, the range between the 25th and 75th percentile), whiskers indicate the 1.5 times of IQR.

**Table 1.** Clinical overview of NSCLC patients treated with ICI.

| Immunotherapy | Targets | No. of samples | No. of patients |
| --- | --- | --- | --- |
| Pembrolizumab | PD-1 | 9 | 8 |
| Atezolizumab | PD-L1 | 1 | 1 |
| Nivolumab | PD-1 | 2 | 1 |
| Vibostolimab+Pembrolizumab | TIGIT + PD-1 | 2 | 1 |

| RECIST | Description | No. of samples | No. of patients |
| --- | --- | --- | --- |
| PR | Partial response | 5 | 4 |
| SD | Stable disease | 3 | 2 |
| PD | Progressive disease | 6 | 5 |

Due to the diversity of sample collection sites, our data may be influenced by varied immune cell composition at different sample collection sites. Therefore, we analyzed immune and stromal cell subsets across early-stage (tLung) and late-stage (tL/B) lung tumors, and metastatic lymph nodes (mLN), comparing them to normal lung (nLung) and lymph node (nLN) tissues. This analysis was conducted on public scRNA-seq data from 43 samples from 33 lung adenocarcinoma (LUAD) patients ^17^ (Fig. S1a-c). Although there were differences in tissue-specific resident populations, we found that the immune cell profiles, especially T/NK cells of mLN were similar to those of primary tumor tissues (Fig. S1d-g).

### Classification of immune cell subset in lung cancer

Global cell-type profiling (Fig. 1b,c and Table S2) illustrates the cellular composition of each sample as epithelial/tumor cells, fibroblasts, endothelial cells, T/natural killer (NK) cells, B/plasma cells, myeloid immune cells, and mast cells. Individual samples show variations in epithelial/tumor content as well as in immune cell composition (Fig. 1d,e). For further analysis of immune cell subtypes, we applied sequential subclustering on global immune cell clusters. As scRNA-seq shows limited performance in separating CD4+ and CD8+ T cell subsets, antibody-derived tag (ADT) information ^18^ was used to complement the transcriptome data and to predict CD4+ T cells, CD8+ T cells, and NK cells (Fig. 2a). Finally, fourteen CD4+ and fourteen CD8+ T cell subclusters were identified excluding <5% ambiguous cells (Fig. 2b,c and Table S2). In the CD4+ T cell compartment, naïve-like T cells (TN, CD4_cluster0) and central memory T cells (TCM, CD4_cluster1) expressing *SELL, TCF7, LEF1,* and *CCR7* genes or tissue-resident memory T cells (TRM, CD4_clusters3, 5, 6) expressing *NR4A1, MYADM,* and *PTGER4* genes were abundant in most samples. Regulatory T cells (Treg, CD4_cluster2) with *FOXP3, CTLA4, ICOS,* and *BATF* expression were also abundant, which has been demonstrated as tumor-specific alterations in the tissue microenvironment ^17,19,20^. In the CD8+ T cell compartment, effector memory T cells (TEM), effector T cells (TEFF), effector memory CD45RA positive cells (TEMRA) (CD8_clusters0, 2, 3, 8) expressing *PRF1* and *IFNG* were dominant over TN/TCM (CD8_clusters4, 5) types. Exhausted T cells (TEX, CD8_clusters1, 12) expressed multiple checkpoint genes (*HAVCR2* and *PDCD1*) along with high levels of *PRF1*, *IFNG*, *CXCR3,* and *CXCL13*. Co-expression of cytotoxic effectors and checkpoint molecules in TEX clusters indicates that cluster populations may retain functional capacity as cytotoxic effector T cells ^21^. Further, clonotype analysis of TCR supported the T cell subset classification demonstrating higher clonal expansion in the CD8+ T cell compartment than that in CD4+ T cells, with the highest levels within the TEX subclass (Fig. S2a-c).

**Figure 2.**
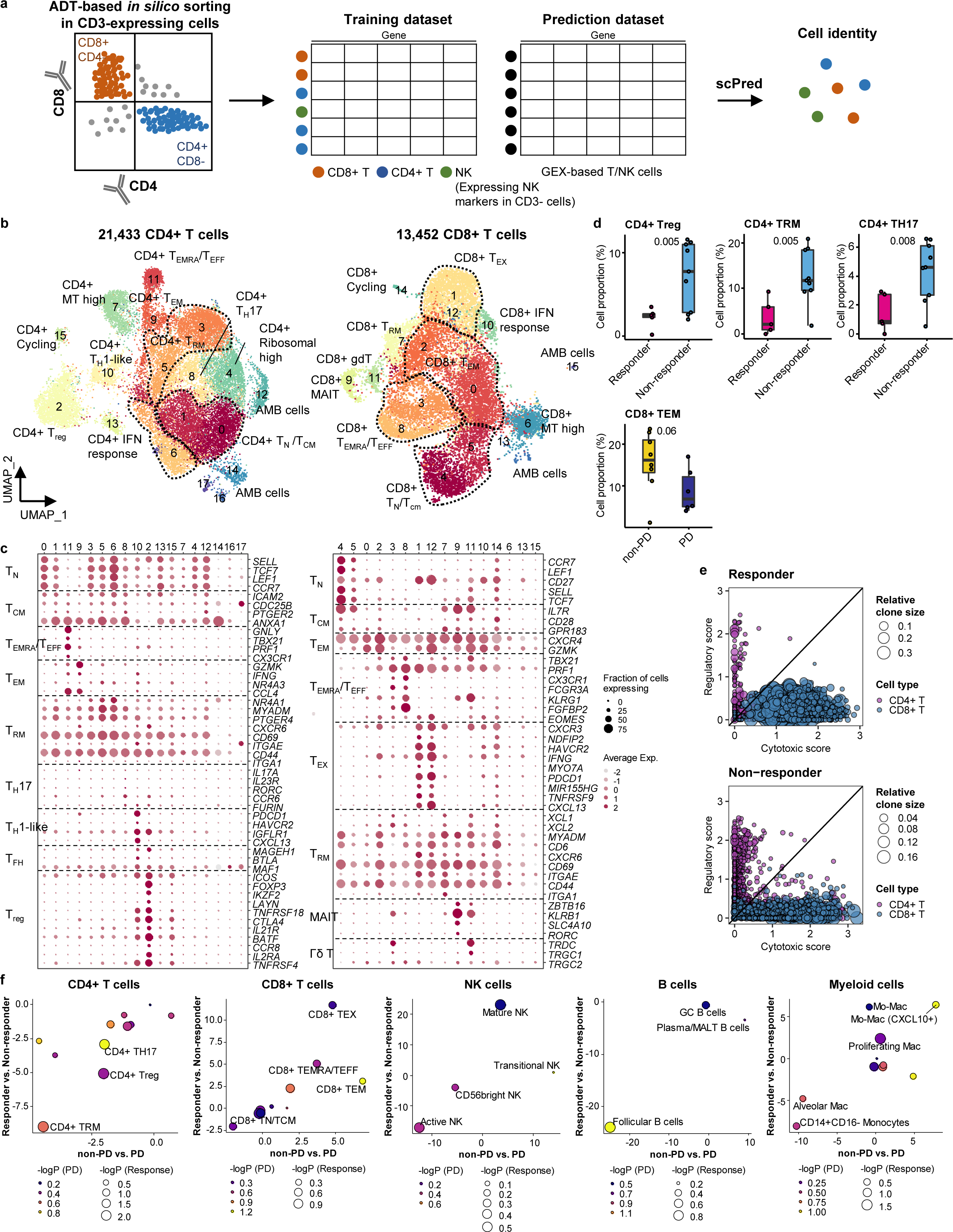
Classification and characterization of CD4+ and CD8+ T cell subtypes. **a,** Prediction strategy to classify CD4+, CD8+ T, and NK cells by applying ADT data from lung cancer. **b,** UMAP plot of CD4+ and CD8+ T cells, colored by clusters. **c,** Dot plot of mean expression of selected CD4+ (left) and CD8+ (right) T cell marker genes in each cell cluster. **d,** Box plot of the percentage of CD4+ and CD8+ T cell types within total CD4+ plus CD8+ cells for sample groups representing responses to ICI. Label represents p-value calculated via two-tailed Student’s t-test. Each box represents the median and the IQR, whiskers indicate the 1.5 times of IQR. **e,** Association of T cell functional features with clonal expansion. Dot size depicts the relative clone size of each cell, which is divided by the total number of CD4+ and CD8+ T cells, respectively. Color indicates the cell lineage. **f,** Comparisons of proportional changes in cell subtypes along ICI responses within each immune cell lineage. The quantitative values shown on the axis represent the mean difference in % cell proportions between sample groups. Dot size and color represent -log (p-value) for responder vs. non-responder and non-PD vs. PD, respectively. The lower left quadrant shows cell types overrepresented in the poor responder groups, while the upper right quadrant indicates cell types overrepresented in the better responder groups. p-value, two-tailed Student’s t-test.

Natural killer cells can be subclassified as CD56bright, transitional, active, and mature types ^22^ (Fig. S3a,b and Table S2). Active NK cells expressed the highest level of *PRF1*, *TNF,* and *IFNG,* reflecting a cytotoxic effector function.

Compared to T cell clusters, fewer B/plasma cells were detected as follicular (B_clusters 0, 1, 2, 4, 6, 11), germinal center (B_cluster 10), and plasma/mucosa-associated lymphoid tissue (MALT) B cells (B_clusters 3, 7, 12) (Fig. S3a,b and Table S2). Plasma/MALT B cells manifested higher levels of clonal expansion of BCR than follicular B cells (Fig. S3c,d). Identical clonality of some follicular B and plasma cells suggests in situ maturation and differentiation of B cells to plasma cells in tumor tissues (Fig. S3c, clonotype 16).

Myeloid cells were composed of monocytes, dendritic cells, and a large number of macrophages (Fig. S3a,b and Table S2). CD14+CD16-classical monocytes (Myeloid_clusters1,6) were predominantly found over non-classical CD14^lo^CD16+ types (Myeloid_cluster15). Alveolar macrophages (Alveolar Mac, Myeloid_cluster0) expressed well-defined marker genes such as *MARCO*, *FABP4*, and *MCEMP1* along with anti-inflammatory genes such as *CD163, APOE,* and *C1QA/B/C*. Monocyte-derived macrophages represent heterogeneous populations with a similar gene expression profile to alveolar macrophages (Mo-Mac, Myeloid_clusters2, 3, 5, 9, 11, 13), with an elevated chemokine gene expression (CXCL10+ Mo-Mac, Myeloid_clusters4, 14, 16), or with active cell cycle progression (Proliferating Mac, Myeloid_cluster7). Dendritic cells were categorized as CD1c+ (Myeloid_cluster8, *CD1C* and *ITGAX*), activated (Myeloid_cluster17, *CCR7* and *LAMP3*), and CD141+ (Myeloid_cluster19, *CLEC9A* and *XCR1*) subclusters. Overall immune cell composition is comparable to those reported in previous studies ^17,19,20^.

### Immune cell landscape fostering the ICI response

In global cell-type profiling (Fig. 1b,c), abundance in T, NK, or myeloid cell types shows no difference between responders and non-responders (Fig. 1d, e). Nonetheless, as specific differentiation features within the cell types may influence the response to ICI treatment, we compared the response groups using the proportion of subclusters within CD4+ T, CD8+ T, NK, B/plasma, and myeloid cells. After subclustering (Fig. 2b and Fig. S3a), three subsets of CD4+ T cells, i.e., Treg, TRM, and CD4+ T helper 17 (TH17), were significantly (p<0.01) overrepresented in the non-responder group (Fig. 2d). In contrast, among CD8+ T cell populations, TEM subsets demonstrated a modest level of association with the patients who responded well against progressive disease (Fig. 2d, p=0.06). TCR clonotype analysis supported the cellular dynamics such that clonal expansion was more prominent in cytotoxic CD8+ T cells over CD4+ Tregs in the responder group (Fig. 2e and Fig. S2c). Overall landscape in each cell type (Fig. 2f) suggests that CD4+ Treg and TRM as well as follicular B cells may interfere with the ICI response, whereas CD8+ T cell activation (TEM, TEMRA/TEFF, and TEX), mature NK cells, and CXCL10+ Mo-Mac cells support the ICI response. The balance between separate immune cell types informs immune regulatory axes that may be targeted to favor the activation of tumor-reactive immunity.

### Systemic evaluation of the immune microenvironment associated with ICI response

Next, we evaluated the immune microenvironment as an entity by using all immune cells as a denominator in the subtype proportions. In this setting, we used diverse clinical group comparisons and identified the immune cell blocks separated by clinical outcomes (Fig. 3a). The immune cell blocks overrepresented in the non-responder groups consisted of CD4+ Treg, follicular B cells, and CD4+ TH17/TRM/ T helper 1 (TH1)-like cells. In the immune cell blocks of the responder groups, CD8+ TEM cells showed the strongest enrichment along with the other CD8+ TEX/TEMRA/TEFF/Mitochondria (MT) high cells as well as CXCL10+ Mo-Mac. Despite the immune footprints of the ICI responders, extensive variations among individual patients (Fig. 3b) hamper patient stratification solely based on the immune profiles.

**Figure 3.**
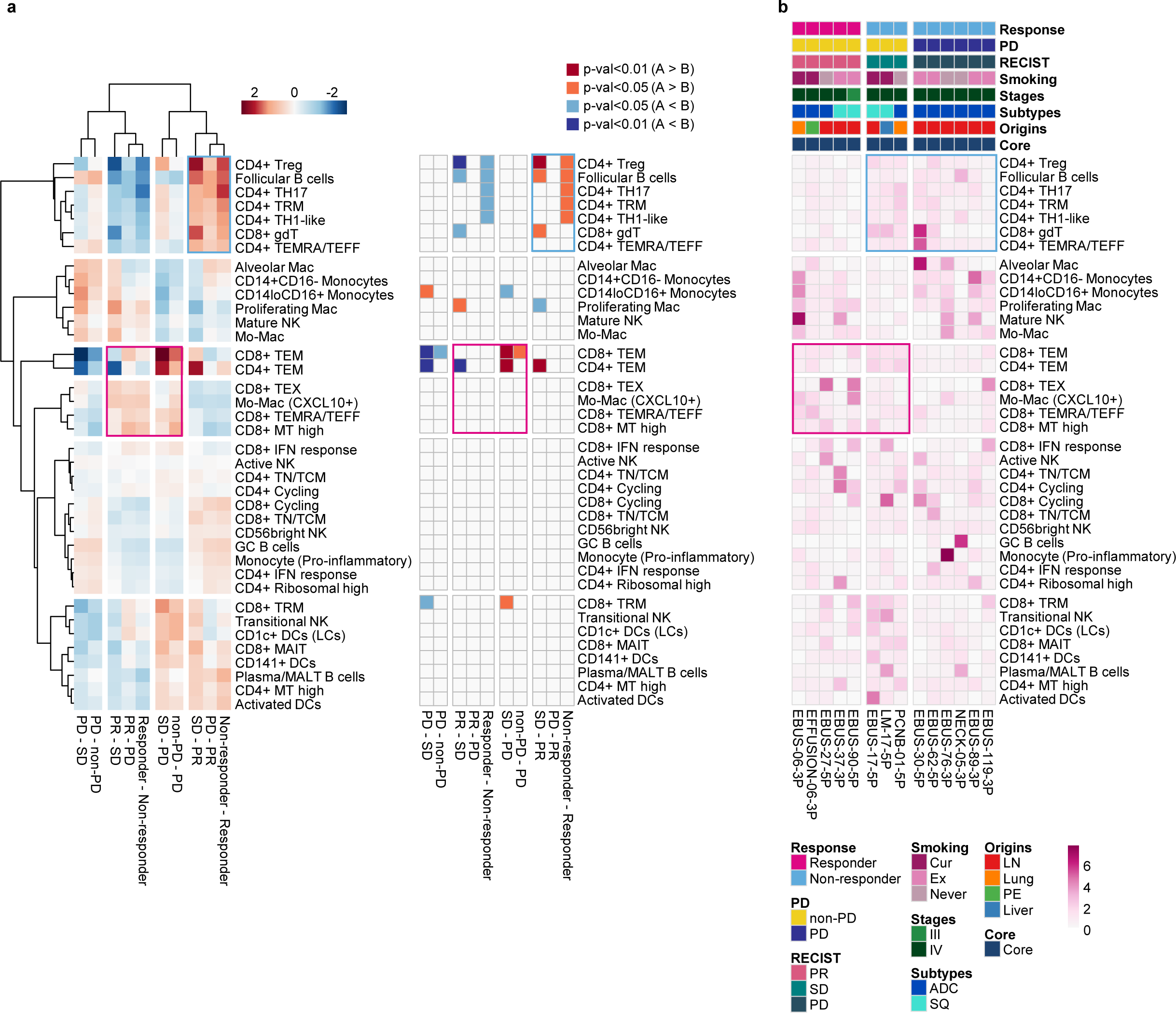
Systemic evaluation of immune cell dynamics associated with response to ICI. **a,** Heat map with unsupervised hierarchical clustering (left) and depicting significance (right) of proportional changes in cell subtypes within total immune cells. Proportional changes were compared for multiple ICI response groups. Color represents the -log (p-value) determined using two-tailed Student’s t-test. **b,** Distribution map for each cell type across individual samples aligned with clinical data. Color represents *Ro/e* score calculated using the chi-square test.

### Tumor cell signatures associated with ICI response

We investigated the associations between genomic characteristics in tumor and ICI response. The constraints of mutation analysis with 10x chromium data complicated the direct correlation between tumor mutation burden and ICI outcome. Rather, we assessed copy number alterations (CNA) indirectly, via chromosomal gene expression patterns. These analyses revealed a moderate correlation between low levels of CNA, including both gain and loss of heterozygosity, and positive responses to ICI (Fig. S4). This finding is consistent with the result from previous genetic association studies ^23^.

To assess gene expression characteristics of tumors influencing ICI response, we separated malignant tumor cell clusters from normal epithelial cell types (Fig. S5). Subsequent DEG analysis (Fig. 4a and Table S3) identified genes in poor response groups linked to the regulation of cell death, cell motility, and cell activation (Fig. S6 and Table S4). The DEGs were refined later by combinations of various tumor signatures separating responder and non-responder groups (Table S5).

**Figure 4.**
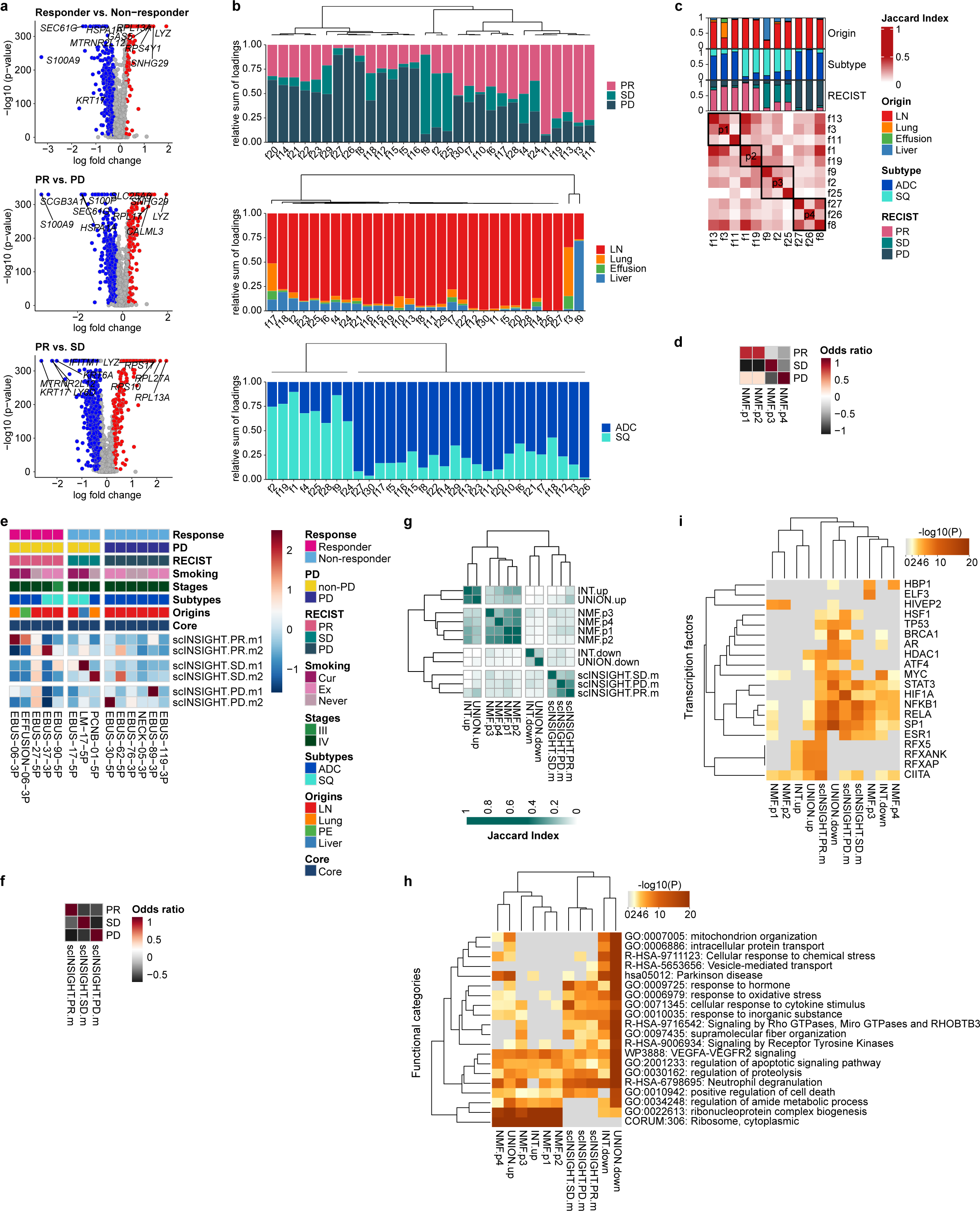
Single-cell tumor signatures associated with response to ICI. **a,** Volcano plot of expression difference for responder vs. non-responder, PR vs. PD, and PR vs. SD in 12,975 malignant cells from 11 core patients. The log fold change indicates the difference in the mean expression level for each gene. The significance level was determined using two-tailed Wilcoxon Rank Sum test. **b,** Relative sum of loadings for all NMF factors contributed to malignant cells from 11 core patients across RECIST, tissue origins, and cancer subtypes, respectively. **c,** Selection of RECIST-enriched NMF programs. **d,** Enrichment of NMF programs for RECIST groups. Color represents the z-transformed odds ratio. **e,** Expression map of RECIST-specific scINSIGHT modules across individual samples aligned with clinical data. Color represents the z-transformed mean expression of genes contributing to each module. **f,** Enrichment of RECIST-specific gene modules for RECIST groups. Color represents the z-transformed odds ratio. **g,** Hierarchical clustering of pairwise similarities between tumor signatures. INT and UNION, intersection and union of DEGs for responder vs. non-responder, PR vs. PD, and PR vs. SD in Fig. 4a. **h,** Functional categories and **i,** transcription factors of the selected tumor signatures, analyzed by Metascape.

Next, to explore the existence of gene programs and modules influencing the ICI response, we applied factorization using non-negative matrix factorization (NMF) and scINSIGHT ^24^. Among 30 factors from NMF across all malignant cells, we identified factors showing high loadings for a specific RECIST group as NMF programs p1∼4 (Fig. 4b, c). There were clear distinction among RECIST groups according to the gene expression levels associated with these NMF programs (Fig. 4d and Table S5). To identify gene modules consistent across different patients, we examined RECIST-specific modules though scINSIGHT analysis ^24^ (Fig. 4e). Unfortunately, we found that contributions to these gene modules varied significantly among patients. To mitigate this variability, we adjusted the RECIST-specific modules by combining genes from the original modules (Table S5). The refined gene modules showed a specific gene expression pattern for each RECIST group, similar to the NMF programs (Fig. 4f). Overall, both genes and their functional categories segregated depending on the selection techniques used (Fig. 4g, h). However, transcription factors governing the signatures derived from DEG, NMF, and scINSIGHT analyses consistently delineated between responders and non-responders (Fig. 4i). Responder-specific gene signatures showed associations with the transcription factors Regulatory Factor X Associated Ankyrin Containing Protein (RFXANK), Regulatory Factor X Associated Protein (RFXAP), and Regulatory Factor X5 (RFX5). This RFX protein complex has emerged as a positive biomarker for the immune response in diverse cancer types ^25^. Non-responder-specific gene signatures were regulated by Activator Of Transcription 3 (STAT3) and Nuclear Factor Kappa B Subunit 1 (NFKB1), known to play roles in PD-L1 regulation and T cell activation in cancer ^26^.

We also adopted principal component analysis (PCA) to isolate correlated gene signatures variably expressed in tumor cells (Fig. S7, and Table S4, S5). Among the top 10 PCs, negatively correlated genes in PC2, PC7, and PC8 distinguished the tumor cells in the poor response groups, whereas positively correlated genes in PC6 and PC9 were upregulated in the better response groups (Fig. S8a-c). Tumor cells from the PR group had low PC2.neg scores, suggesting low growth factor/type I interferon response signaling as a tumor cell-specific positive predictor of the ICI response. Conversely, high levels of growth factor/type I interferon response signaling in tumor cells may present intrinsic resistance to PD-(L)1 inhibitor alone or in combination. Type I interferon is known to drive anti-tumor effect directly or indirectly on tumor and surrounding immune cells, but also acts to counter the anti-tumor effect by inducing CD8+ T cell exhaustion and up-regulating immune-suppressive genes on tumor cells ^27^. We assessed whether tumor cell signatures are applicable in association with ICI response in other tumors. They had a modest influence on the response to ICI treatment of melanoma (Fig. S8d) in bulk gene expression data ^28,29^.

These tumor cell signatures were specifically linked to elucidating the ICI response, but did not demonstrate any prognostic value for LUAD or lung squamous cell carcinoma (LUSC) in the TCGA RNA sequencing data (Table 2).

**Table 2.** Multivariate overall survival analysis of tumor signature genes.

|  | TCGA LUAD |  |  |  | TCGA LUSC |  |  |  |
| --- | --- | --- | --- | --- | --- | --- | --- | --- |
|  | HR | Lower CI | Upper CI | p-value | HR | Lower CI | Upper CI | p-value |
| NMF.p1 | 0.89 | 0.53 | 1.48 | 0.64 | 0.92 | 0.66 | 1.27 | 0.60 |
| NMF.p2 | 0.87 | 0.53 | 1.43 | 0.59 | 0.93 | 0.67 | 1.28 | 0.66 |
| NMF.p3 | 0.76 | 0.49 | 1.21 | 0.25 | 1.06 | 0.76 | 1.49 | 0.72 |
| NMF.p4 | 1.00 | 0.64 | 1.55 | 0.98 | 1.00 | 0.70 | 1.44 | 0.98 |
| scINSIGHT.PR.m | 0.97 | 0.63 | 1.51 | 0.90 | 0.87 | 0.62 | 1.23 | 0.44 |
| scINSIGHT.SD.m | 0.81 | 0.53 | 1.23 | 0.32 | 0.94 | 0.67 | 1.33 | 0.74 |
| scINSIGHT.PD.m | 0.89 | 0.57 | 1.40 | 0.62 | 0.87 | 0.62 | 1.24 | 0.45 |
| PC1.pos | 0.48 | 0.27 | 0.84 | 0.01 | 0.78 | 0.55 | 1.12 | 0.17 |
| PC1.neg | 0.98 | 0.65 | 1.48 | 0.92 | 0.72 | 0.49 | 1.06 | 0.10 |
| PC2.pos | 0.87 | 0.55 | 1.36 | 0.53 | 0.98 | 0.69 | 1.40 | 0.91 |
| PC2.neg | 0.82 | 0.53 | 1.25 | 0.35 | 0.90 | 0.63 | 1.27 | 0.54 |
| PC3.pos | 1.14 | 0.68 | 1.90 | 0.63 | 0.87 | 0.63 | 1.19 | 0.38 |
| PC3.neg | 0.86 | 0.55 | 1.35 | 0.52 | 0.79 | 0.54 | 1.16 | 0.23 |
| PC4.pos | 0.88 | 0.54 | 1.42 | 0.59 | 0.66 | 0.46 | 0.95 | 0.02 |
| PC4.neg | 1.10 | 0.75 | 1.62 | 0.62 | 0.97 | 0.69 | 1.35 | 0.84 |
| PC5.pos | 1.09 | 0.71 | 1.68 | 0.68 | 0.76 | 0.53 | 1.10 | 0.15 |
| PC5.neg | 1.01 | 0.69 | 1.50 | 0.95 | 1.03 | 0.71 | 1.49 | 0.88 |
| PC6.pos | 0.77 | 0.45 | 1.29 | 0.32 | 1.09 | 0.79 | 1.51 | 0.60 |
| PC6.neg | 0.68 | 0.41 | 1.13 | 0.13 | 0.81 | 0.58 | 1.13 | 0.21 |
| PC7.pos | 1.10 | 0.71 | 1.72 | 0.67 | 0.75 | 0.51 | 1.10 | 0.14 |
| PC7.neg | 0.88 | 0.58 | 1.33 | 0.53 | 0.77 | 0.55 | 1.09 | 0.14 |
| PC8.pos | 0.77 | 0.50 | 1.18 | 0.23 | 1.01 | 0.72 | 1.43 | 0.94 |
| PC8.neg | 1.00 | 0.65 | 1.52 | 0.99 | 0.95 | 0.66 | 1.37 | 0.80 |
| PC9.pos | 0.91 | 0.60 | 1.38 | 0.66 | 1.02 | 0.73 | 1.43 | 0.91 |
| PC9.neg | 1.11 | 0.73 | 1.68 | 0.63 | 1.02 | 0.72 | 1.45 | 0.89 |
| PC10.pos | 1.13 | 0.76 | 1.68 | 0.55 | 1.28 | 0.88 | 1.86 | 0.20 |
| PC10.neg | 0.93 | 0.59 | 1.46 | 0.75 | 0.93 | 0.65 | 1.35 | 0.71 |
| INT.up | 0.82 | 0.53 | 1.26 | 0.37 | 1.00 | 0.71 | 1.42 | 0.99 |
| UNION.up | 0.80 | 0.52 | 1.21 | 0.28 | 0.77 | 0.53 | 1.13 | 0.18 |
| INT.down | 0.84 | 0.56 | 1.24 | 0.37 | 1.07 | 0.73 | 1.57 | 0.74 |
| UNION.down | 0.67 | 0.42 | 1.06 | 0.08 | 0.75 | 0.52 | 1.07 | 0.11 |

### Combination of tumor signatures and immune cell dynamics classify ICI response

Immune cell dynamics or tumor signature alone has a limited capacity to profile the therapeutic outcome of PD-(L)1 inhibitor alone or in combination (Fig. 5a). Similar to the immune cell blocks and tumor signatures that were over-represented in the poor response group, each of CD4+ Treg, CD4+ TH17, PC7.neg, INT.down, and UNION.down was significantly associated with ICI response in univariate regression analysis (Fig. 5b). PC7.neg denotes genes negatively correlated with PC7, a principal component extracted from PCA that distinguishes tumor cells in poor response groups. INT.down and UNION.down represent the intersection and union of down-regulated genes in the responder group, respectively. The variation in ICI response was not affected by clinical variables of tissue origin, cancer subtype, pathological stage, and smoking status. When we performed a combined analysis of the top tumor-immune features to classify response, the discriminative power (AUC) was improved to over 95% (Fig. 5c). Overall, features of the non-responders, especially CD4+ Treg, B/Plasma cells, INT.down, and UNION.down, showed a higher estimate than those of responders. These non-responder features suggest heterogeneous mechanisms of resistance conferred by tumor and immune regulatory axes.

**Figure 5.**
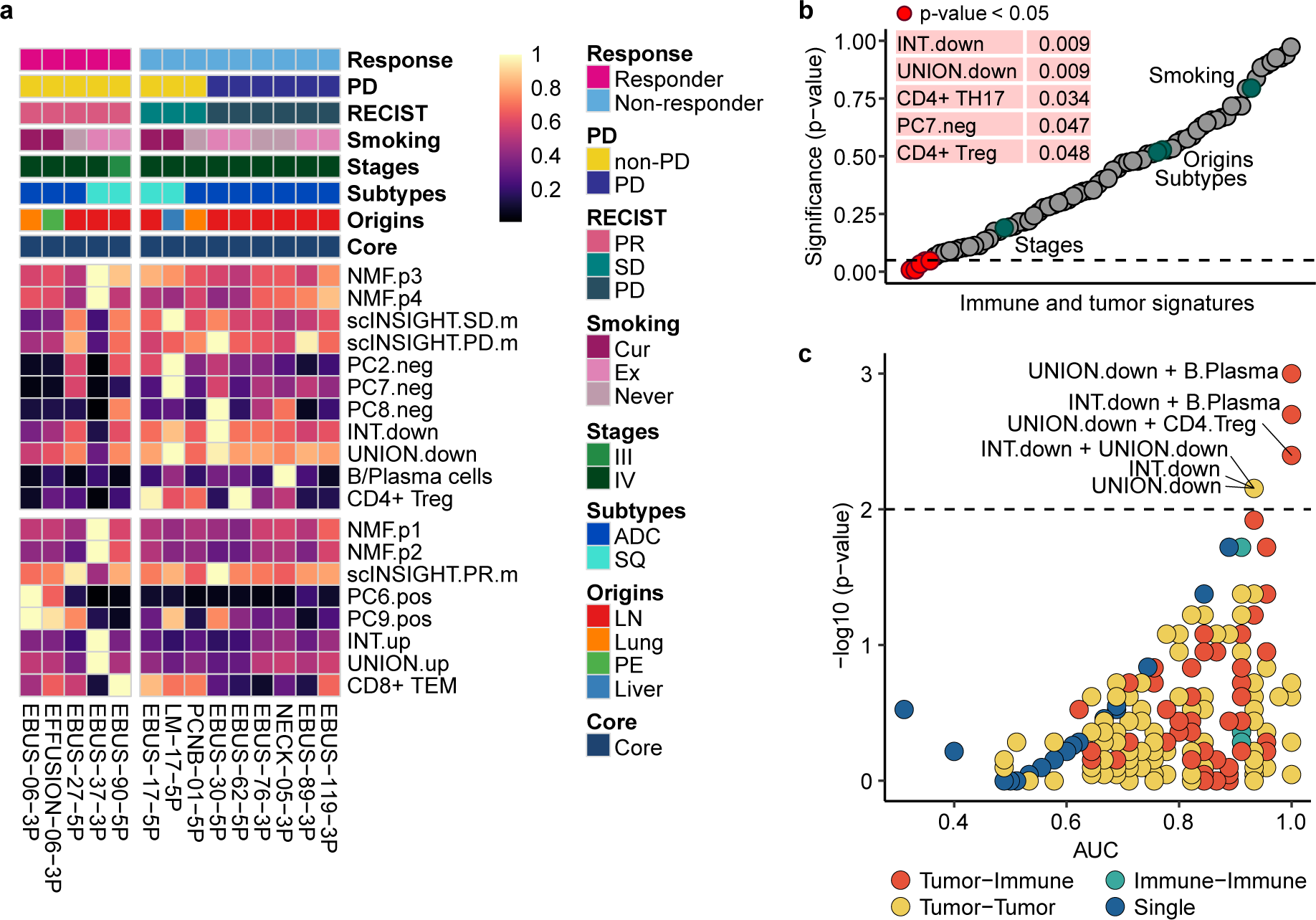
Combination of tumor signatures and immune index classifying the response to ICI. **a,** Heat map of relative contribution of tumor signatures and immune index across individual samples aligned with clinical data. Mean expression of each tumor signature and the percentage of each immune index are divided by the maximum value across samples. INT and UNION, intersection and union of DEGs for responder vs. non-responder, PR vs. PD, and PR vs. SD in Fig. 4a. **b,** Univariate regression analysis of immune and tumor signatures for ICI response, together with clinical variables. **c,** ROC analysis of combinatorial index to classify responder and non-responder. p-value, two-tailed Wilcoxon Rank Sum test. P-values adjusted with the Benjamini-Hochberg correction; UNION.down + B/Plasma, 0.22; INT.down + B/Plasma, 0.22; UNION.down + CD4+ Treg, 0.26; INT.down + UNION.down, 0.26; INT.down, 0.26; UNION.down, 0.26.

## Discussion

ICI alone, or in combination with chemotherapy, are considered standard first-line therapy for patients with NSCLC. NSCLC may harbor large numbers of genetic perturbations due to genotoxic environmental exposure, which likely generate high mutation burden or neoantigens ^30^. The neoantigen-directed T cell response is hampered by diverse immune suppressive mechanisms exerted by tumor cells and the immune regulatory network ^31^. Current ICIs targeting PD-(L)1 aim one angle of many suppressive mechanisms, and identifying the features of non-responders will reveal additional regulatory angles to improve ICI response.

Immune regulatory network would determine the balance between activation and suppression of tumor-directed immunity. In previous studies, prediction of the response to ICI highlighted CD8+ cytotoxic effector T cells and CD4+ Tregs ^32^. The involvement of effector CD8+ T cells is consistent in most studies regardless of the cellular origin (blood or tissues) or tumor type (melanoma, lung cancer) ^33^. Their phenotypes slightly differ depending on the cellular resolution of the study. Our data provided the highest resolution cell types, and CD8+ TEM cells (*GZMK*, *CXCR4* expression) were overrepresented in the responders. By comparison, Tregs were underrepresented in the responder group. Previously, our group reported a contrasting result demonstrating an increase in Tregs in the posttreatment blood samples (not baseline) in the responder group ^34^. This discrepancy can be explained by the differences in the measurements, sites, and timing of sampling, and the resolution of subpopulations. In mouse preclinical models, PD-L1 inhibitor treatment induces T cell expansion of all phenotypes including CD4+/CD8+ TEFF and Tregs ^35^. Thus, Treg expansion captured in the posttreatment blood samples may represent overall immune activation in human patients. Alternatively, heterogeneity of Tregs and complex effects of PD-(L)1 inhibition on this cell type may contribute to variable results in response prediction.

The identification of an abundance of CD4+ TRM cells as a negative predictor of ICI response is an unexpected finding, considering that higher frequencies of TRM cells in lung tumor tissues are generally associated with better clinical outcomes in NSCLC ^36^. This is largely due to their role in sustaining high densities of tumor-infiltrating lymphocytes and promoting anti-tumor responses. Additionally, previous studies have demonstrated that TRM cell subsets coexpressing PD-1 and TIM-3 are relatively enriched in patients who respond to PD-1 inhibitors ^37^. However, recent findings suggest that pre-existing TRM-like cells in lung cancer may promote immune evasion mechanisms, contributing to resistance to immune checkpoint blockade therapies ^38^. These observations suggest that the roles of TRM subsets in tumor immunity are highly context-dependent.

Similarly, CD4+ TH17 cells, which were overrepresented in the non-responder groups, exhibit context-dependent roles in tumor immunity and may be associated with both unfavorable and favorable outcomes ^39,40^. In exploring tumor cell signatures linked to ICI response, non-responder attributes were regulated by STAT3 and NFKB1. The STAT3 and NF-κB pathways are crucial for Th17 cell differentiation and T cell activation ^41,42^. Notably, STAT3 activation in lung cancer orchestrates immunosuppressive characteristics by inhibiting T-cell mediated cytotoxicity ^43^. The combined influence of the Th17/STAT3 axis and TRM cell activity in predicting ICI response underscores the complexity of these pathways and suggests that their roles in tumor immunity and therapy response warrants further investigation.

In search of tumor cell signatures associated with the response to ICI, we adopted two approaches, an individual gene level comparison between the responder and non-responder group and a feature extraction approach to decompose data using the NMF and PCA. Both approaches highlighted attributes of non-responders governed by key transcription factors, which play a significant role in immune response regulation. The ability to predict ICI response based on tumor signatures was as accurate as predictions based on immune cell behavior. Integrating data from both immune and tumor cells enhanced the discriminative power (AUC) for identifying responder, suggesting the presence of both interactive and distinct mechanisms of resistance.

Our study has limitations. Primarily, most samples were obtained from metastatic lymph nodes rather than original tumor tissues, potentially not reflecting the tumor microenvironment accurately. However, prior study ^17^ and Fig. S1 have shown that the immune microenvironment within metastatic lymph nodes closely resembles that of lung tumor tissues, rather than normal lymph nodes. On a positive note, our findings indicate that the immune landscape of metastatic lymph nodes can predict ICI response. Another challenge is the small sample size and the issue of gene expression drop-out, necessitating further studies with a larger patient cohort. Despite these limitations, our study stands out by employing high-throughput scRNA-seq to both tumor cells and immune microenvironment, offering a comprehensive analysis of the multicellular factors that affect ICI response in patients with advanced NSCLC.

## Materials and Methods

### Human specimens

This study was approved by the Institutional Review Board (IRB) of Samsung Medical Center (IRB no. 2010-04-039-052). Informed written consent was obtained from all patients enrolled in the study. The study participants included 26 patients diagnosed with lung cancer (Table S1). The study population (n=26) has been treated with the investigator’s choice either as a clinical trial (n=5) or as standard clinical practice (n=21). Regardless of the treatment selection, the specimens were prospectively collected based on the study protocol. A total of 33 samples were collected and immediately transferred on ice for tissue preparation. Metastatic lymph nodes, metastatic liver tissues, and lung/bronchus tumor tissues from patients with lung cancer were collected using endobronchial ultrasound bronchoscopy, neck lymph node ultrasound and biopsy, liver biopsy, and percutaneous transthoracic cutting needle biopsy. Tumors, normal lungs, normal lymph nodes, and normal brain tissues were obtained during resection surgery. Pleural fluid was collected from patients with malignant pleural effusion.

### Clinical outcomes

The clinical outcomes of ICI were evaluated based on the Response Evaluation Criteria in Solid Tumor (RECIST) 1.1 ^44^. In this study, we described non-responders as patients with a stable or progressive disease. Responders were considered as patients with a partial response. None of the patients showed a complete response.

### Sample preparation

Single-cell isolation was performed differently depending on the samples. (1) Biopsy samples and normal lymph node tissues were chopped into 2-4 mm pieces and dissociated in an enzyme solution containing collagenase/hyaluronidase (STEMCELL Technologies, Vancouver, Canada) and DNase I, RNase-Free (lyophilized) (QIAGEN, Hilden, Germany) at 37°C for 1 h. Tissue pieces were re-mixed by gentle pipetting at 20-min intervals during incubation. (2) Tumor and normal lung tissue dissociation was performed using a tumor dissociation kit (Miltenyi Biotech, Germany) following the manufacturer’s instructions. Briefly, tissue was cut into 2–4 mm pieces and transferred to a C tube containing the enzyme mix (enzymes H, R, and A in RPMI1640 medium). The GentleMACS programs h_tumor_01, h_tumor_02, and h_tumor_02 were run with two 30-min incubations on a MACSmix tube rotator at 37°C. (3) Brain tissue was chopped into 2–4 mm pieces and incubated in an enzyme solution (collagenase (Gibco, Waltham, MA, USA), DNase I (Roche, Basel, Switzerland), and Dispase I (Gibco) in DMEM) at 37°C for 1 h. Tissue pieces were re-mixed by gentle pipetting at 15-min intervals during incubation. (4) Pleural fluids were transferred to a 50-ml tube, and the cells were spun down at 300g.

Each cell suspension was transferred to a new 50-ml (15-ml for biopsy samples) tube through a 70-µm strainer. The volume in the tube was readjusted to 50-ml (or 15-ml) with RPMI1640 medium, and spun down to remove the enzymes. The supernatant was aspirated, the cell pellet was resuspended in 4 ml of RPMI1640 medium, and dead cells were removed using Ficoll-Paque PLUS (GE Healthcare, Chicago, IL, USA) separation.

For samples subjected to multiplexing, dissociated cells were cryopreserved in CELLBANKER1 (Zenogen, Fukushima, Japan) and thawed for pooling.

### Single-cell RNA sequencing (scRNA-seq) and read processing

Single-cell suspensions were loaded into a Chromium system (10x Genomics, Pleasanton, CA, USA). Following the manufacturer’s instructions, 3’ scRNA-seq libraries for the 14 samples were generated using Chromium Single Cell 3′ v2 Reagent Kits. The 3’ library preparation for EBUS_119 used Chromium Single Cell 3′ v3 Reagent Kits. The 5’ scRNA-seq libraries for twelve individual and two pooled samples were generated using Chromium Single Cell 5′ v2 Reagent Kit. Libraries were then sequenced on an Illumina HiSeq 2500 for 3’ scRNA-seq and an Illumina NovaSeq 6000 for 5’ scRNA-seq. Sequencing reads were mapped to the GRCh38 human reference genome using Cell Ranger toolkit (v5.0.0).

### SNP genotyping array

Genomic DNA was extracted from the peripheral blood of six patients and subjected to sample multiplexing (DNeasy Blood & Tissue Kit, QIAGEN). The 766,221 single nucleotide polymorphisms (SNPs) was genotyped using Illumina Global Screening Array MG v2, following the manufacturer’s instructions. Normalized signal intensity and genotype were processed using Illumina’s GenomeStudio v.2 software.

### Demultiplexing of pooled samples

The individuals in sample multiplexing were assigned by a software tool *freemuxlet*, which is an extension of *demuxlet* ^45^ (https://github.com/statgen/popscle). First, the popscle tool *dsc-pileup* was run with the bam file generated by Cell Ranger toolkit and reference vcf file. The reference was assembled after a lift-over process with GRCh38 from 1000 Genomes Project phase 1 data and the variant allele frequency in East Asian >0.01 were discarded. Next, *freemuxlet* was used to determine the sample identity with default parameters. The individuals were matched based on the similarity between freemuxlet-annotated genotypes and SNP array-detected genotypes.

### Acquisition of scRNA-seq data from lung adenocarcinoma (LUAD) patients

We obtained raw 3’ scRNA-seq from 43 specimens acquired from 33 LUAD patients including early-stage (tLung) and late-stage (tL/B) lung tumor tissues, metastatic lymph nodes (mLN), normal lung tissues (nLung) and lymph nodes (nLN) ^17^. Sequencing reads were mapped to the GRCh38 human reference genome using Cell Ranger toolkit (v5.0.0).

### scRNA-seq data analysis

The raw gene-cell-barcode matrix from Cell Ranger pipeline was processed using Seurat v3.2.2 R package ^46^. Cells were selected using two quality criteria: mitochondrial genes (<20%) and gene count (>200). Cell multiplets predicted by Scrublet ^47^ were filtered out. From the filtered cells, the unique molecular identifier (UMI) count matrix was log-normalized and scaled by z-transform while regressing out the effects of cell-cycle variations for subsequent analysis. For batch correction, we used Harmony v1.0 R package ^48^ interfacing with Seurat as the *RunHarmony* function. A total of 2,000 variably expressed genes were selected using *FindVariableFeatures* with a parameter selection.method=“vst”. A subset of principal components (PCs) was selected based on *ElbowPlot* function. Uniform Manifold Approximation and Projection (UMAP) for dimension reduction and cell clustering was performed using *RunUMAP, FindNeighbors*, and *FindClusters* functions with the selected PCs and resolutions [Advanced lung cancer patients (Total cells, 33 PCs and resolution=0.3; CD4+ T cells, 28 PCs and resolution=0.9; CD8+ T cells, 28 PCs and resolution=0.9; NK cells, 26 PCs and resolution=0.3; B/Plasma cells, 28 PCs and resolution=0.3; Myeloid cells, 30 PCs and resolution=1.2), LUAD patients (Total cells, 23 PCs and resolution=0.3; T/NK cells, 24 PCs and resolution=0.9)]. We applied the *FindAllMarkers* function to identify differentially expressed genes (DEGs) for each cell cluster. Significance was determined using Wilcoxon Rank Sum test. Genes were selected according to the following statistical thresholds; log fold change>0.25, p-value<0.01, adjusted p-value (bonferroni correction)<0.01, and percentage of cells (pct)>0.25. Cell identity was determined by comparing the expression of known marker genes and DEGs for each cluster.

### Principal component analysis (PCA) analysis using the proportion of cell lineages and T/NK cell subsets

PCA analysis was performed for the % proportion of cell lineages and T/NK cell subsets in individual LUAD samples using *prcomp* function of stats v3.6.3 R package. For total cells, the percentages of immune and stromal cells were calculated except for epithelial, cycling, and AMB (ambiguous) cells. For T/NK cells, unknown cells annotated as MT high and AMB cells were excluded.

### In silico classification of CD4+ T, CD8+ T, and NK cells

We characterized CD4+ T, CD8+ T, and NK cell populations by combined analysis of gene and protein expression using Cellular Indexing of Transcriptomes and Epitopes by Sequencing data from primary tumor and normal lungs. Among the cells in clusters annotated as T/NK cells, we identified CD3-expressing cells with CD3D or CD3E or CD3G >0 at the RNA level. CD4 and CD8 positive cells were then identified with a cutoff at 55th percentile of antibody-derived tags (ADT) level. NK cells were identified based on the RNA expression level of NK cell markers (*XCL1*, *NCAM1*, *KLRD1*, and *KLRF1*) in CD3 negative cells. The gene expression matrix with cell identity of CD4 positive, CD8 positive, and NK was applied as reference data for supervised cell-type classification using *getFeatureSpace* and *trainModel* functions of scPred v1.9.0 R package ^49^. Finally, we classified T/NK cells in Fig. 1b into CD4+ T, CD8+ T, and NK cells using scPred *scPredict* function.

### Analysis of TCR/BCR repertoires in CD4+ T, CD8+ T, and B/Plasma cells

The data derived from Cell Ranger pipeline for T cell receptor (TCR) and B cell receptor (BCR) sequencing data were processed using scRepertoire v1.2.0 R package ^50^ in R v4.1.1. We selected contigs that generated alpha-beta chain pairs for TCR and heavy-light chain pairs for BCR for subsequent analysis. We called clonotypes based on V(D)JC genes and CDR3 nucleotide sequence with the parameter clonecall=“gene+nt”. The set of clone types was classified by total frequency using the parameter cloneTypes defined as Single=1, Small=5, Medium=10, Large=20, and Hyperexpanded=Inf.

### Scoring of T cell functional features

Scores for T cell functional features were calculated as the mean expression of regulatory (*ICOS*, *FOXP3*, *IKZF2*, *LAYN*, *TNFRSF18*, *CTLA4*, *IL21R*, *BATF*, *CCR8*, *IL2RA*, and *TNFRSF4*) and cytotoxic (*CX3CR1*, *PRF1*, *GZMA*, *GZMB*, *GZMH*, *GNLY*, *KLRG1*, and *NKG7*) genes at the log-normalized level.

### Identification of malignant cells based on inferred copy number variation (CNV) from scRNA-seq data

Two computational tools, inferCNV v1.2.1 (https://github.com/broadinstitute/inferCNV) and CopyKAT v1.0.5 R packages ^51^, were used to infer genomic copy numbers from scRNA-seq. In a run with inferCNV, the UMI count matrix of each tumor sample was loaded into inferCNV *CreateInfercnvObject* function along with cell lineage annotations. The reference (normal) cells were selected as cells annotated with T/NK, B/Plasma, myeloid, and mast cells. We maintained the proportion of epithelial cells below 20% in each tumor sample using the expression profiles of the normal lung and lymph node tissues. Inferred CNV signals were analyzed using inferCNV *run* function using the parameters: cutoff=0.1, denoise=TRUE, HMM=TRUE, and HMM_type=“i6”. The signals were then summarized as standard deviations (s.d.) for all windows and the correlation between the CNV in each cell and the mean of the top 5% cells ^52^. Cancer cells showing CNV perturbation (>0.03 s.d. or >0.3 CNV correlation) were classified as malignant cells, otherwise as non-malignant cells. The UMI count matrix of each tumor sample was loaded into CopyKAT *copykat* function along with cell lineage annotations using the following parameters: ngene.chr=3, KS.cut=0.05, and norm.cell.names. Cancer cells predicted as aneuploid cells by CopyKAT were classified as malignant cells. Finally, we identified malignant cells, which are cancer cells classified as malignant cells in either inferCNV or CopyKAT.

### Single-cell DEGs between response groups in malignant cells

A total of 12,975 malignant cells were used to identify DEGs in pairwise comparisons according to responder versus non-responder, PR versus PD, and PR versus SD. Differential expression levels were calculated using Seurat *FindMarkers* function with the Wilcoxon Rank Sum test. Genes were selected according to the following statistical thresholds: log fold change>0.25, p-value<0.01, adjusted p-value (bonferroni correction)<0.01, and pct>0.25. We reconstructed DEGs as four groups: INT.up, INT.down, UNION,up, and UNION.down, based on with the intersection (INT) and union (UNION) of up- or down-regulated genes for pairwise comparisons between responder versus non-responder, PR versus PD, and PR versus SD. INT.up and INT.down represent the intersection of up- and down-regulated genes in the responder group, respectively. UNION.up and UNION.down represent the union of up- and down-regulated genes in the responder group, respectively.

### Non-negative matrix factorization (NMF) programs of the malignant cells

The UMI count matrix for malignant cells was loaded into *nmf* function of RcppML v0.5.6 R package. A NMF model was learned with a rank of 30 using all genes. For each of the 30 NMF factors, the top-ranked 50 genes in the NMF score were defined as signatures. RECIST-enriched NMF program consisted of selected factors based on their relative sum of loadings. We aggregated and redefined gene signatures of factors included in each NMF program. The uniqueness of each NMF program for RECIST groups was evaluated as an odds ratio using *fisher.test* function of stats v3.6.3 R package. Annotations of NMF programs were assigned using Metascape ^53^.

### RECIST-specific gene modules in malignant cells

The RECIST-specific gene modules were analyzed with a matrix factorization named scINSIGHT ^24^ using log-normalized count and 2,000 highly variable genes for each sample. For each module, we selected the 100 genes with the highest coefficients. Combinations of the top 100 genes for modules specific to each RECIST group were defined as module genes. The uniqueness of each module for RECIST groups was evaluated as an odds ratio using *fisher.test* function of stats v3.6.3 R package. Annotations of gene modules were assigned using Metascape ^53^.

### Principal component signatures of the malignant cells

The UMI count matrix for malignant cells was log-normalized and scaled by z-transform while regressing out the effects of cell-cycle variations for PCA. A total of 2,000 variably expressed genes selected using *FindVariableFeatures* with selection.method=“vst” were used for PCA. PCs were calculated by Seurat *RunPCA* function. PC signatures were selected for 30 genes with + (pos) and – (neg) scores that highly contributed to each PC from PC1 to PC10.

### Functional category analysis

Functional categories representing the enriched gene expression in comparisons for responder vs. non-responder, PR vs. PD, and PR vs. SD as well as in the PCs were identified using fgsea v1.12.0 ^54^ R package with parameters: minSize=10, maxSize=600, and nperm=10000. Gene sets for Gene Ontology (GO) Biological Process were collected from the MSigDB database using msigdbr v7.1.1 R package ^55,56^. The gene list was ranked by the log fold change for each comparison and feature loadings for each PC. Significant GO terms were selected after collapsing redundant terms using fgsea *collapsePathways* function with a statistical threshold for Benjamini-Hochberg adjusted p-value<0.05.

### Multivariate overall survival analysis of tumor signatures

To evaluate the prognostic potential of tumor signatures, RNA sequencing data from LUAD and LUSC samples were retrieved from The Cancer Genome Atlas (TCGA) data portal (https://portal.gdc.cancer.gov/) ^57^. The dataset comprised 533 primary tumors from TCGA LUAD and 502 primary tumors from TCGA LUSC. Gene expression levels were quantified as (log2 FPKM-UQ + 1), where FPKM-UQ represents the upper quartile fragments per kilobase per million mapped reads for each sample.

Clinical variables were first categorized, including age (below and above median age), gender (male and female), pathological stage (I/II and III/IV), distant metastasis (M0 and M1), and nodal status (N0 and Ns). Samples were then classified into high and low groups based on the upper quantile of the mean expression for each tumor signature group. Multivariate survival analysis was conducted using the *analyse_multivariate* function of the survivalAnalysis R package.

### Evaluation of discriminative power of identifying responders for tumor signatures and combinatorial indexes

Classification models of responders and non-responders for PC signatures and combinatorial indexes between tumor and/or immune cells were generated based on in-sample performance and tested by Receiver Operating Characteristic (ROC) curve. Relative numbers between the observed and expected cells (*Ro/e*) for each sample were obtained from the chi-square test ^19^. To describe the separability, area under the curve (AUC) was calculated using *ROC* function of Epi v2.44 R package with *Ro/e* scores as input. Significance was calculated by Wilcoxon Rank Sum test and confirmed by adjusting with the Benjamini-Hochberg correction.

### Univariate regression analysis for ICI response

Univariate regression was performed using the *lm* function of stats v3.6.3 with *Ro/e* scores for each sample as input. We evaluated the relationship between the target variable ICI response, classified as responders and non-responders, and one predictor variable of immune cell types, tumor signatures, and clinical factors such as tissue origin, cancer subtype, pathological stage, and smoking status. The significance of predictor was calculated using *Anova* function of car v3.0-9 R package.

### Validation of PC signatures in melanoma cohorts

We used Riaz et al.’s ^29^ and Van Allen al.’s ^28^ RNA sequencing data from melanoma patients receiving PD-1 and CTLA-4 immune checkpoint therapy to assess expressional changes of PC signatures along RECIST. The mean expression of each PC signature in each RECIST group was calculated as the log2 normalized level.

### Data availability

Raw single-cell RNA sequencing data generated during the current study are available in the European Genome-phenome Archive (EGA) database (accession code EGAD00001008703), and processed data can be accessed from the NCBI Gene Expression Omnibus (GEO) database (accession code GSE205335).

Single-cell RNA sequencing data for LUAD patients analyzed in this study are available in the EGA database at EGAD00001005054. Bulk RNA sequencing data analyzed in this study were obtained from GEO at GSE91061 and database of Genotypes and Phenotypes (dbGap) at phs000452.v2.p1.

## Acknowledgements

This study was supported by the Collaborative Genome Program for Fostering New Post-Genome Industry (NRF-2017M3C9A6044633 and NRF-2017M3C9A6044636), Mid-Career Researcher Program (NRF-2022R1A2C1091451), and Basic Research Laboratory Program (RS-2023-00220840) of the National Research Foundation of Korea funded by the Korea government. We also acknowledge the Basic Medical Science Facilitation Program, through the Catholic Medical Center of the Catholic University of Korea funded by the Catholic Education Foundation and the KREONET/GLORIAD service provided by KISTI (Korea Institute of Science and Technology Information).

## Supplementary Files

**Supplementary file 1: Fig. S1-S8. Supplementary figures and legends.**

**Supplementary file 2: Table S1. Patient information of lung cancer ICI cohorts.**

**Supplementary file 3: Table S2. List of genes specific to the cell clusters in total cells and each cell lineage.**

**Supplementary file 4: Table S3. List of DEGs in comparison between ICI response groups.**

**Supplementary file 5: Table S4. Details of GO terms significantly enriched in comparisons for ICI response groups and PCs.**

**Supplementary file 6: Table S5. Gene list of tumor signatures.**

## References

1 Reck, M. et al. Pembrolizumab versus Chemotherapy for PD-L1-Positive Non-Small-Cell Lung Cancer. N Engl J Med 375, 1823–1833, doi:10.1056/NEJMoa1606774 (2016).

2 Gandhi, L. et al. Pembrolizumab plus Chemotherapy in Metastatic Non-Small-Cell Lung Cancer. N Engl J Med 378, 2078–2092, doi:10.1056/NEJMoa1801005 (2018).

3 Paz-Ares, L. et al. Pembrolizumab plus Chemotherapy for Squamous Non-Small-Cell Lung Cancer. N Engl J Med 379, 2040–2051, doi:10.1056/NEJMoa1810865 (2018).

4 Wakelee H, A. N., Zhou C, Csőszi T, Vynnychenko IO, Goloborodko O, Luft A, Andrey Akopov, Martinez-Marti A, Kenmotsu H, Chen Y, Chella A, Sugawara S, Gitlitz BJ, Bennett E, Wu F, Yi J, Deng Y, McCleland M, Felip E. IMpower010: Primary results of a phase III global study of atezolizumab versus best supportive care after adjuvant chemotherapy in resected stage IB-IIIA non-small cell lung cancer (NSCLC). J Clin Oncol 39, 8500, doi:10.1200/JCO.2021.39.15_suppl.8500 (2021).

5 Reck, M. et al. Five-Year Outcomes With Pembrolizumab Versus Chemotherapy for Metastatic Non-Small-Cell Lung Cancer With PD-L1 Tumor Proportion Score >/= 50. J Clin Oncol 39, 2339–2349, doi:10.1200/JCO.21.00174 (2021).

6 Fehrenbacher, L. et al. Atezolizumab versus docetaxel for patients with previously treated non-small-cell lung cancer (POPLAR): a multicentre, open-label, phase 2 randomised controlled trial. Lancet 387, 1837–1846, doi:10.1016/S0140-6736(16)00587-0 (2016).

7 Hellmann, M. D. et al. Nivolumab plus Ipilimumab in Lung Cancer with a High Tumor Mutational Burden. N Engl J Med 378, 2093–2104, doi:10.1056/NEJMoa1801946 (2018).

8 Litchfield, K. et al. Meta-analysis of tumor- and T cell-intrinsic mechanisms of sensitization to checkpoint inhibition. Cell 184, 596–614 e514, doi:10.1016/j.cell.2021.01.002 (2021).

9 Kamphorst, A. O. et al. Proliferation of PD-1+ CD8 T cells in peripheral blood after PD-1-targeted therapy in lung cancer patients. Proc Natl Acad Sci U S A 114, 4993–4998, doi:10.1073/pnas.1705327114 (2017).

10 Kim, K. H. et al. The First-week Proliferative Response of Peripheral Blood PD-1(+)CD8(+) T Cells Predicts the Response to Anti-PD-1 Therapy in Solid Tumors. Clin Cancer Res 25, 2144–2154, doi:10.1158/1078-0432.CCR-18-1449 (2019).

11 Thommen, D. S. et al. A transcriptionally and functionally distinct PD-1(+) CD8(+) T cell pool with predictive potential in non-small-cell lung cancer treated with PD-1 blockade. Nat Med 24, 994–1004, doi:10.1038/s41591-018-0057-z (2018).

12 Simon, S. & Labarriere, N. PD-1 expression on tumor-specific T cells: Friend or foe for immunotherapy? Oncoimmunology 7, e1364828, doi:10.1080/2162402X.2017.1364828 (2017).

13 Arce Vargas, F., et al. Fc-Optimized Anti-CD25 Depletes Tumor-Infiltrating Regulatory T Cells and Synergizes with PD-1 Blockade to Eradicate Established Tumors. Immunity 46, 577–586, doi:10.1016/j.immuni.2017.03.013 (2017).

14 Krieg, C. et al. Author Correction: High-dimensional single-cell analysis predicts response to anti-PD-1 immunotherapy. Nat Med 24, 1773–1775, doi:10.1038/s41591-018-0094-7 (2018).

15 Kumagai, S. et al. The PD-1 expression balance between effector and regulatory T cells predicts the clinical efficacy of PD-1 blockade therapies. Nat Immunol 21, 1346–1358, doi:10.1038/s41590-020-0769-3 (2020).

16 Ruiz-Patino, A. et al. Immunotherapy at any line of treatment improves survival in patients with advanced metastatic non-small cell lung cancer (NSCLC) compared with chemotherapy (Quijote-CLICaP). Thorac Cancer 11, 353–361, doi:10.1111/1759-7714.13272 (2020).

17 Kim, N. et al. Single-cell RNA sequencing demonstrates the molecular and cellular reprogramming of metastatic lung adenocarcinoma. Nat Commun 11, 2285, doi:10.1038/s41467-020-16164-1 (2020).

18 Stoeckius, M. et al. Simultaneous epitope and transcriptome measurement in single cells. Nat Methods 14, 865–868, doi:10.1038/nmeth.4380 (2017).

19 Guo, X. et al. Global characterization of T cells in non-small-cell lung cancer by single-cell sequencing. Nat Med 24, 978–985, doi:10.1038/s41591-018-0045-3 (2018).

20 Gueguen, P. et al. Contribution of resident and circulating precursors to tumor-infiltrating CD8(+) T cell populations in lung cancer. Sci Immunol 6, doi:10.1126/sciimmunol.abd5778 (2021).

21 Groom, J. R. & Luster, A. D. CXCR3 in T cell function. Exp Cell Res 317, 620–631, doi:10.1016/j.yexcr.2010.12.017 (2011).

22 Yang, C. et al. Heterogeneity of human bone marrow and blood natural killer cells defined by single-cell transcriptome. Nat Commun 10, 3931, doi:10.1038/s41467-019-11947-7 (2019).

23 Liu, L. et al. Combination of TMB and CNA Stratifies Prognostic and Predictive Responses to Immunotherapy Across Metastatic Cancer. Clin Cancer Res 25, 7413–7423, doi:10.1158/1078-0432.CCR-19-0558 (2019).

24 Qian, K., Fu, S., Li, H. & Li, W. V. scINSIGHT for interpreting single-cell gene expression from biologically heterogeneous data. Genome Biol 23, 82, doi:10.1186/s13059-022-02649-3 (2022).

25 Lapuente-Santana, O., van Genderen, M., Hilbers, P. A. J., Finotello, F. & Eduati, F. Interpretable systems biomarkers predict response to immune-checkpoint inhibitors. Patterns (N Y) 2, 100293, doi:10.1016/j.patter.2021.100293 (2021).

26 Betzler, A. C. et al. NF-kappaB and Its Role in Checkpoint Control. Int J Mol Sci 21, doi:10.3390/ijms21113949 (2020).

27 Fenton, S. E., Saleiro, D. & Platanias, L. C. Type I and II Interferons in the Anti-Tumor Immune Response. Cancers (Basel) 13, doi:10.3390/cancers13051037 (2021).

28 Van Allen, E. M. et al. Genomic correlates of response to CTLA-4 blockade in metastatic melanoma. Science 350, 207–211, doi:10.1126/science.aad0095 (2015).

29 Riaz, N. et al. Tumor and Microenvironment Evolution during Immunotherapy with Nivolumab. Cell 171, 934–949 e916, doi:10.1016/j.cell.2017.09.028 (2017).

30 Lawrence, M. S. et al. Mutational heterogeneity in cancer and the search for new cancer-associated genes. Nature 499, 214–218, doi:10.1038/nature12213 (2013).

31 Sharma, P., Hu-Lieskovan, S., Wargo, J. A. & Ribas, A. Primary, Adaptive, and Acquired Resistance to Cancer Immunotherapy. Cell 168, 707–723, doi:10.1016/j.cell.2017.01.017 (2017).

32 Gibellini, L. et al. Single-Cell Approaches to Profile the Response to Immune Checkpoint Inhibitors. Front Immunol 11, 490, doi:10.3389/fimmu.2020.00490 (2020).

33 Zheng, L. et al. Pan-cancer single-cell landscape of tumor-infiltrating T cells. Science 374, abe6474, doi:10.1126/science.abe6474 (2021).

34 Koh, J. et al. Regulatory (FoxP3(+)) T cells and TGF-beta predict the response to anti-PD-1 immunotherapy in patients with non-small cell lung cancer. Sci Rep 10, 18994, doi:10.1038/s41598-020-76130-1 (2020).

35 Wei, S. C. et al. Combination anti-CTLA-4 plus anti-PD-1 checkpoint blockade utilizes cellular mechanisms partially distinct from monotherapies. Proc Natl Acad Sci U S A 116, 22699–22709, doi:10.1073/pnas.1821218116 (2019).

36 Ganesan, A. P. et al. Tissue-resident memory features are linked to the magnitude of cytotoxic T cell responses in human lung cancer. Nat Immunol 18, 940–950, doi:10.1038/ni.3775 (2017).

37 Clarke, J. et al. Single-cell transcriptomic analysis of tissue-resident memory T cells in human lung cancer. J Exp Med 216, 2128–2149, doi:10.1084/jem.20190249 (2019).

38 Weeden, C. E. et al. Early immune pressure initiated by tissue-resident memory T cells sculpts tumor evolution in non-small cell lung cancer. Cancer Cell 41, 837–852 e836, doi:10.1016/j.ccell.2023.03.019 (2023).

39 Marques, H. S. et al. Relationship between Th17 immune response and cancer. World J Clin Oncol 12, 845–867, doi:10.5306/wjco.v12.i10.845 (2021).

40 Chang, S. H. T helper 17 (Th17) cells and interleukin-17 (IL-17) in cancer. Arch Pharm Res 42, 549–559, doi:10.1007/s12272-019-01146-9 (2019).

41 Park, S. H., Cho, G. & Park, S. G. NF-kappaB Activation in T Helper 17 Cell Differentiation. Immune Netw 14, 14–20, doi:10.4110/in.2014.14.1.14 (2014).

42 Poholek, C. H. et al. Noncanonical STAT3 activity sustains pathogenic Th17 proliferation and cytokine response to antigen. J Exp Med 217, doi:10.1084/jem.20191761 (2020).

43 Jing, B. et al. IL6/STAT3 Signaling Orchestrates Premetastatic Niche Formation and Immunosuppressive Traits in Lung. Cancer Res 80, 784–797, doi:10.1158/0008-5472.CAN-19-2013 (2020).

44 Eisenhauer, E. A. et al. New response evaluation criteria in solid tumours: revised RECIST guideline (version 1.1). Eur J Cancer 45, 228–247, doi:10.1016/j.ejca.2008.10.026 (2009).

45 Kang, H. M. et al. Multiplexed droplet single-cell RNA-sequencing using natural genetic variation. Nat Biotechnol 36, 89–94, doi:10.1038/nbt.4042 (2018).

46 Stuart, T. et al. Comprehensive Integration of Single-Cell Data. Cell 177, 1888–1902 e1821, doi:10.1016/j.cell.2019.05.031 (2019).

47 Wolock, S. L., Lopez, R. & Klein, A. M. Scrublet: Computational Identification of Cell Doublets in Single-Cell Transcriptomic Data. Cell Syst 8, 281–291 e289, doi:10.1016/j.cels.2018.11.005 (2019).

48 Korsunsky, I. et al. Fast, sensitive and accurate integration of single-cell data with Harmony. Nat Methods 16, 1289–1296, doi:10.1038/s41592-019-0619-0 (2019).

49 Alquicira-Hernandez, J., Sathe, A., Ji, H. P., Nguyen, Q. & Powell, J. E. scPred: accurate supervised method for cell-type classification from single-cell RNA-seq data. Genome Biol 20, 264, doi:10.1186/s13059-019-1862-5 (2019).

50 Borcherding, N., Bormann, N. L. & Kraus, G. scRepertoire: An R-based toolkit for single-cell immune receptor analysis. F1000Res 9, 47, doi:10.12688/f1000research.22139.2 (2020).

51 Gao, R. et al. Delineating copy number and clonal substructure in human tumors from single-cell transcriptomes. Nat Biotechnol 39, 599–608, doi:10.1038/s41587-020-00795-2 (2021).

52 Puram, S. V. et al. Single-Cell Transcriptomic Analysis of Primary and Metastatic Tumor Ecosystems in Head and Neck Cancer. Cell 171, 1611–1624 e1624, doi:10.1016/j.cell.2017.10.044 (2017).

53. Zhou, Y., et al. Metascape provides a biologist-oriented resource for the analysis of systems-level datasets. Nat Commun 10, 1523, doi:10.1038/s41467-019-09234-6 (2019).

54. Korotkevich G, S. V., Sergushichev A. Fast gene set enrichment analysis. bioRxiv, doi:10.1101/060012 (2019).

55 Liberzon, A. et al. Molecular signatures database (MSigDB) 3.0. Bioinformatics 27, 1739–1740, doi:10.1093/bioinformatics/btr260 (2011).

56 Subramanian, A. et al. Gene set enrichment analysis: a knowledge-based approach for interpreting genome-wide expression profiles. Proc Natl Acad Sci U S A 102, 15545–15550, doi:10.1073/pnas.0506580102 (2005).

57 Grossman, R. L. et al. Toward a Shared Vision for Cancer Genomic Data. N Engl J Med 375, 1109–1112, doi:10.1056/NEJMp1607591 (2016).

