## Supplementary figures for "Unveiling the influence of tumor and immune signatures on immune checkpoint therapy in advanced lung cancer"

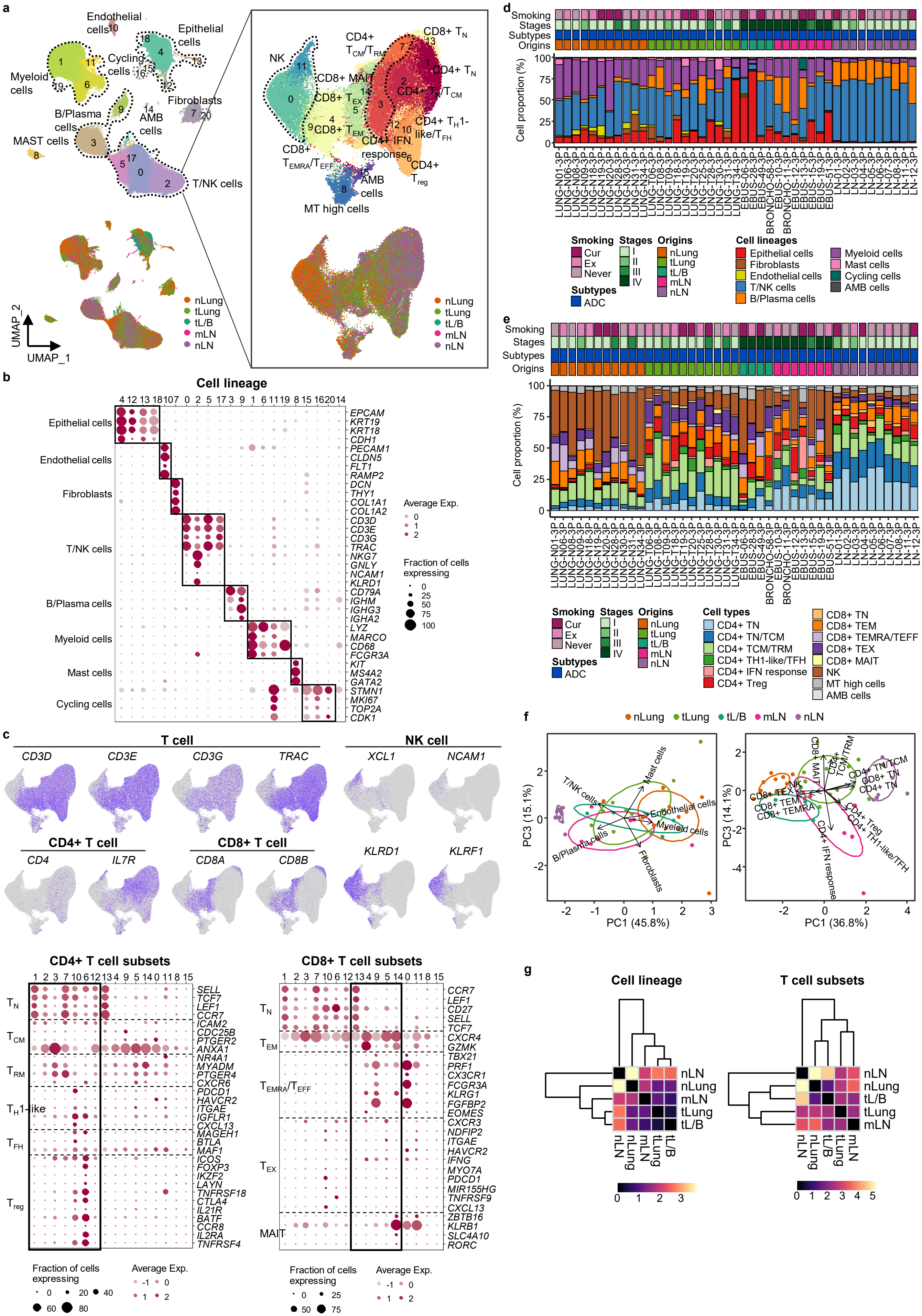

a
Endothelial cells
Epithelial cells
Myeloid cells
Cycling cells
Fibroblasts
B/Plasma cells
AMB cells
MAST cells
T/NK cells
UMAP_2
UMAP_1
nLung
tLung
tL/B
mLN
nLN
NK
CD4+ $T_{CM}/T_{RM}$
CD8+ $T_N$
CD4+ $T_N$
CD8+ MAIT
CD8+ $T_{EX}$
CD4+ $T_N/T_{CM}$
CD8+ $T_{EM}$
CD4+ $T_H$1-like/$T_{FH}$
CD4+ IFN response
CD8+ $T_{EMRA}/T_{EFF}$
CD4+ $T_{reg}$
AMB cells
MT high cells
b
Cell lineage
Epithelial cells
Endothelial cells
Fibroblasts
T/NK cells
B/Plasma cells
Myeloid cells
Mast cells
Cycling cells
EPCAM
KRT19
KRT18
CDH1
PECAM1
CLDN5
FLT1
RAMP2
DCN
THY1
COL1A1
COL1A2
CD3D
CD3E
CD3G
TRAC
NKG7
GNLY
NCAM1
KLRD1
CD79A
IGHM
IGHG3
IGHA2
LYZ
MARCO
CD68
FCGR3A
KIT
MS4A2
GATA2
STMN1
MKI67
TOP2A
CDK1
Average Exp.
Fraction of cells expressing
c
T cell
NK cell
CD3D
CD3E
CD3G
TRAC
XCL1
NCAM1
CD4+ T cell
CD8+ T cell
CD4
IL7R
CD8A
CD8B
KLRD1
KLRF1
CD4+ T cell subsets
CD8+ T cell subsets
$T_N$
$T_{CM}$
$T_{RM}$
$T_H$1-like
$T_{FH}$
$T_{reg}$
$T_{EM}$
$T_{EMRA}/T_{EFF}$
$T_{EX}$
MAIT
SELL
TCF7
LEF1
CCR7
ICAM2
CDC25B
PTGER2
ANXA1
NR4A1
MYADM
PTGER4
CXCR6
PDCD1
HAVCR2
ITGAE
IGFLR1
CXCL13
MAGEH1
BTLA
MAF1
ICOS
FOXP3
IKZF2
LAYN
TNFRSF18
CTLA4
IL21R
BATF
CCR8
IL2RA
TNFRSF4
CD27
CXCR4
GZMK
TBX21
PRF1
CX3CR1
FCGR3A
KLRG1
FGFBP2
EOMES
CXCR3
NDFIP2
IFNG
MYO7A
MIR155HG
TNFRSF9
ZBTB16
KLRB1
SLC4A10
RORC
Fraction of cells expressing
Average Exp.
d
Smoking
Stages
Subtypes
Origins
Cell proportion (%)
Smoking: Cur, Ex, Never
Stages: I, II, III, IV
Origins: nLung, tLung, tL/B, mLN, nLN
Subtypes: ADC
Cell lineages: Epithelial cells, Fibroblasts, Endothelial cells, T/NK cells, B/Plasma cells, Myeloid cells, Mast cells, Cycling cells, AMB cells
e
Cell types: CD4+ TN, CD4+ TN/TCM, CD4+ TCM/TRM, CD4+ TH1-like/TFH, CD4+ IFN response, CD4+ Treg, CD8+ TN, CD8+ TEM, CD8+ TEMRA/TEFF, CD8+ TEX, CD8+ MAIT, NK, MT high cells, AMB cells
f
PC3 (15.1%)
PC1 (45.8%)
PC3 (14.1%)
PC1 (36.8%)
T/NK cells
Mast cells
Endothelial cells
Myeloid cells
B/Plasma cells
Fibroblasts
CD4+ TN/TCM
CD8+ TN
CD4+ TN
CD4+ Treg
CD4+ TH1-like/TFH
CD4+ IFN response
CD8+ TEX
CD8+ TEM
CD8+ TEMRA
g
Cell lineage
T cell subsets
nLN
nLung
mLN
tLung
tL/B

**Fig. S1. Immune cell profiles for sample collection sites in LUAD. a,** UMAP plot of 162,651 single cells (left) and 74,071 T/NK cells (right) from 43 samples acquired from 33 LUAD patients, colored by clusters (top) and tissue origins (bottom). AMB cells, Ambiguous cells. **b,** Dot plot of mean expression of selected marker genes in each cell cluster of 162,651 cells from 43 samples acquired from 33 LUAD patients. **c,** UMAP plot of T/NK cells, colored by expression of T, NK, CD4+ T, and CD8+ T cell marker genes (top). Dot plot of mean expression of selected CD4+ (bottom left) and CD8+ (bottom right) T cell marker genes in each cell cluster. **d, e,** Proportions of the cell lineages (d) and T cell subsets (e) in LUAD tissue from 43 specimens shown by individual samples aligned with clinical data. Cur, Current smoker; Ex, Ex-smoker; Never, Never smoker; ADC, Adenocarcinoma; nLung, normal lung; tLung, early stage tumor lung; tL/B, advanced stage tumor lung; mLN, lymph node metastases; nLN, normal lymph node. **f,** Biplot of PCA for cell lineage (left) and T/NK cell subset (right) proportions. **g,** Euclidean distance map of centroids between groups of samples based on their tissue of origin in the PC1 and PC3 plot in Fig. S1f.

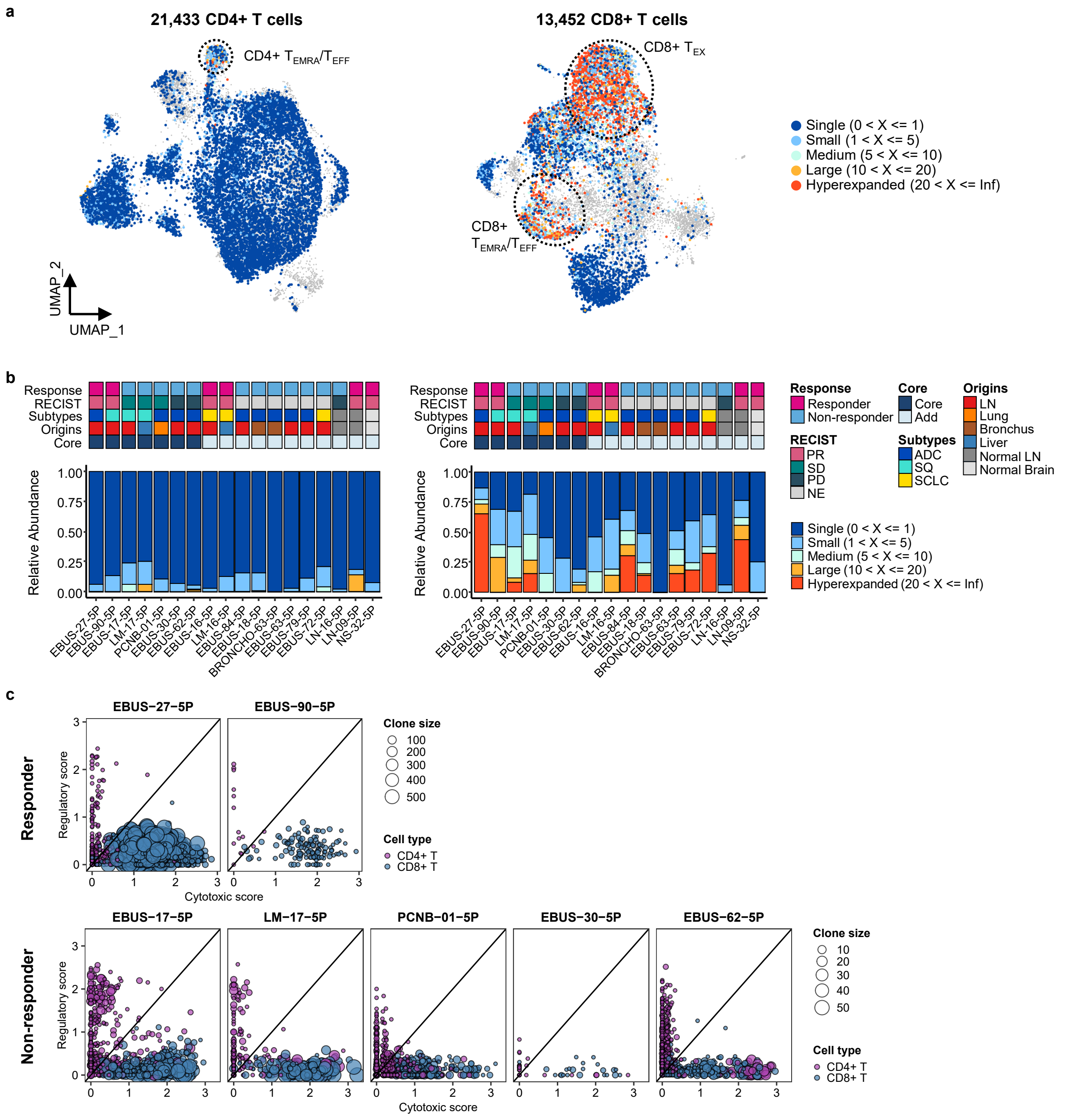


**Fig. S2. Features of TCR repertoires of CD4+ and CD8+ T cells. a,** UMAP plot of CD4+ and CD8+ T cells, colored by clone types (single to hyperexpanded). **b,** Proportions of clone types of CD4+ (left) and CD8+ (right) T cells in individual samples aligned with clinical data. **c,** Association of T cell functional features with clonal expansion in individual samples. Dot size depicts the clone size of each cell and color indicates the cell lineage.

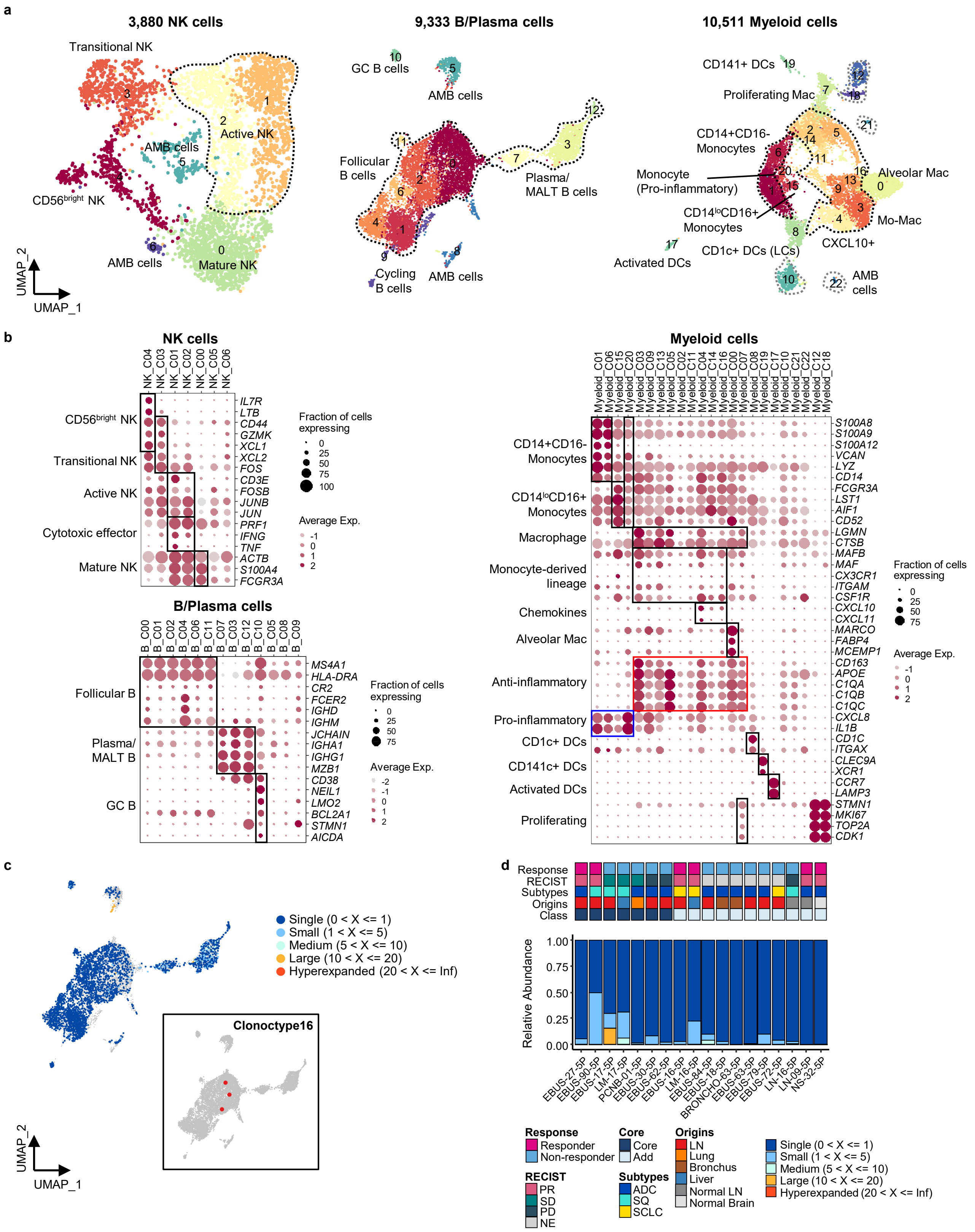


**Fig. S3. Heterogeneity in NK, B, and myeloid cells and features of BCR repertoires. a,** UMAP plot of NK, B, and myeloid cells, colored by clusters. **b,** Dot plot of mean expression of selected marker genes in each cell cluster of NK, B, and myeloid cells. **c,** UMAP plot of B cells, colored by clone types (single to hyperexpanded). Clone distribution of clonotype 16 in B cells (inset). **d,** Proportions of clone types of B cells in individual samples aligned with clinical data.

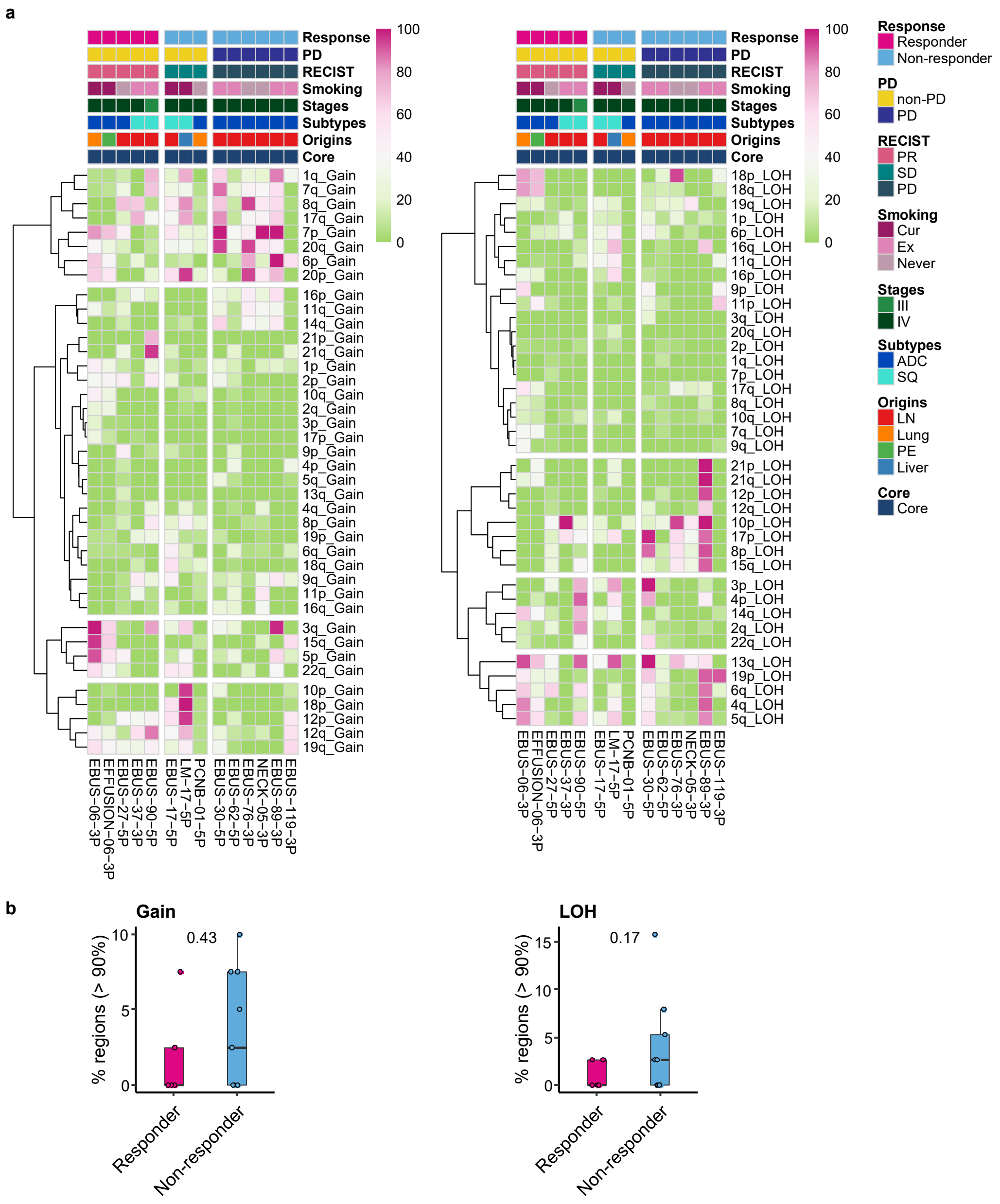


**Fig. S4. CNA profiles in NSCLC tissue aligned with ICI response. a,** Percent map of single-cell CNA events in individual samples aligned with clinical data. The CNV signals of epithelial cells inferred from scRNA-seq were calculated using inferCNV R package. Color represents the percentage of the gain (left) and loss of heterozygosity (LOH) (right) events in individual cells from each sample. **b,** Comparison of % regions showing high perturbation (> 90%) of CNA events between responder and non-responder. Label represents p-value calculated via two-tailed Student's t-test. Each box represents the median and the IQR, whiskers indicate the 1.5 times of IQR.

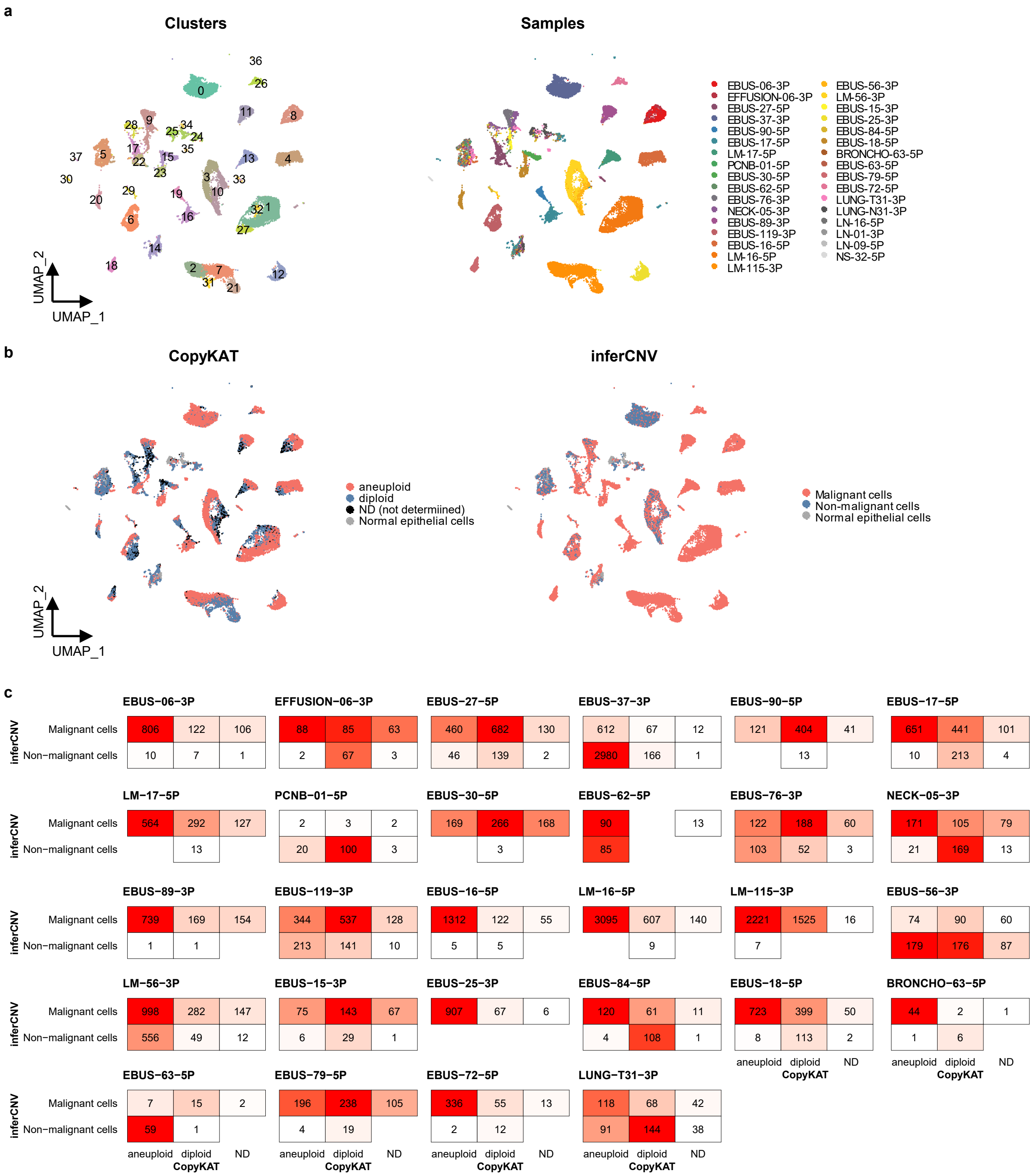


**Fig. S5. Classification of malignant cells based on genomic perturbations. a,** UMAP plot of 31,519 epithelial cells from 26 patients, projected without batch correction, colored by clusters (left) and samples (right). **b,** UMAP projection as shown in Fig. S5a, colored by malignant cell classification predicted using CopyKAT (left) and inferCNV (right) R packages. **c,** Comparisons of malignant cell classification between CopyKAT and inferCNV in each individual sample.

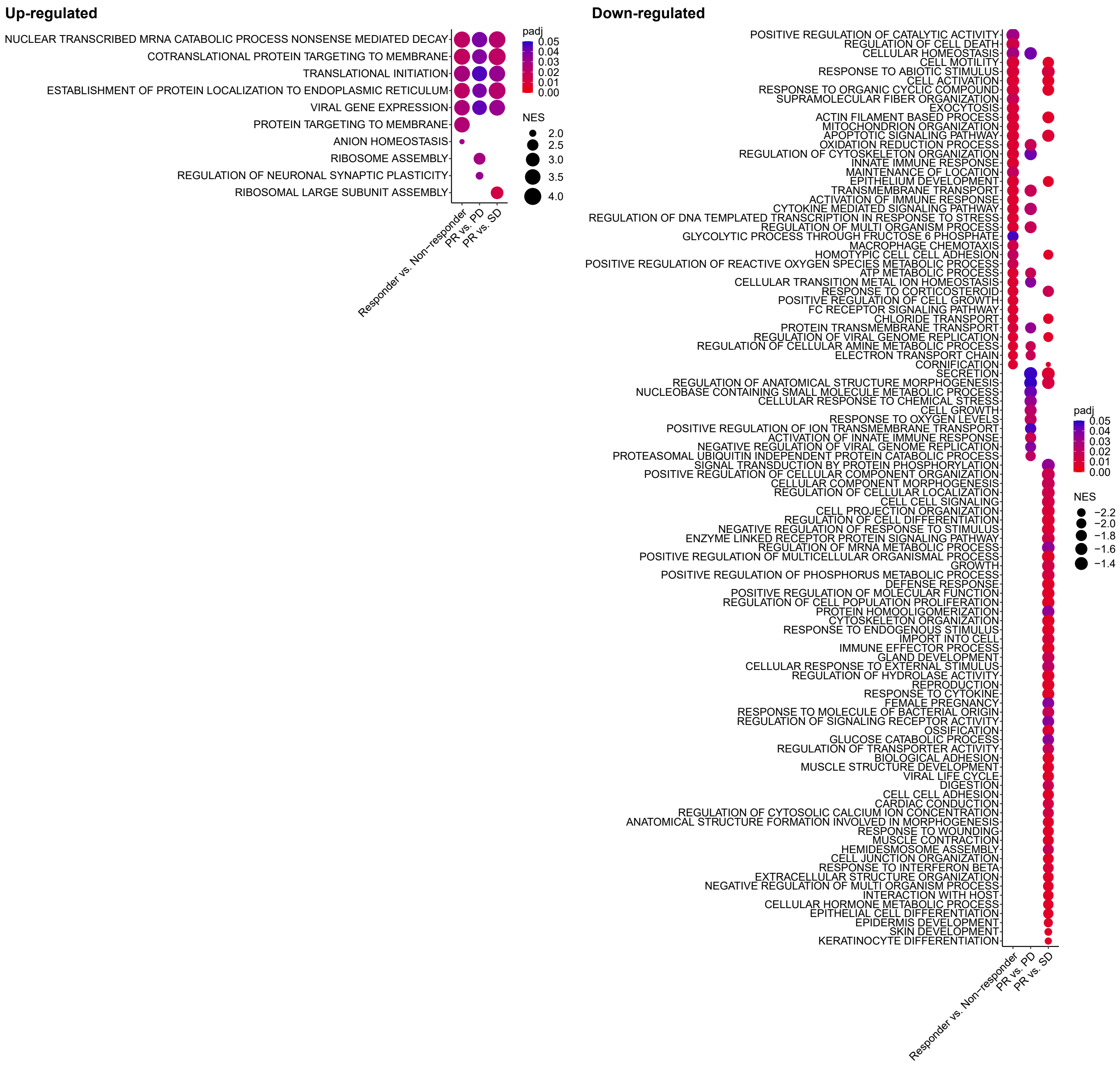


**Fig. S6. Functional categories of single-cell DEG signatures associated with response to ICI.** Enrichment map of significant GO terms for each single-cell DEG signature associated with response to ICI. Color and size indicate the adjusted p-value (padj) and the normalized enrichment score (NES) calculated using fgsea R package, respectively.

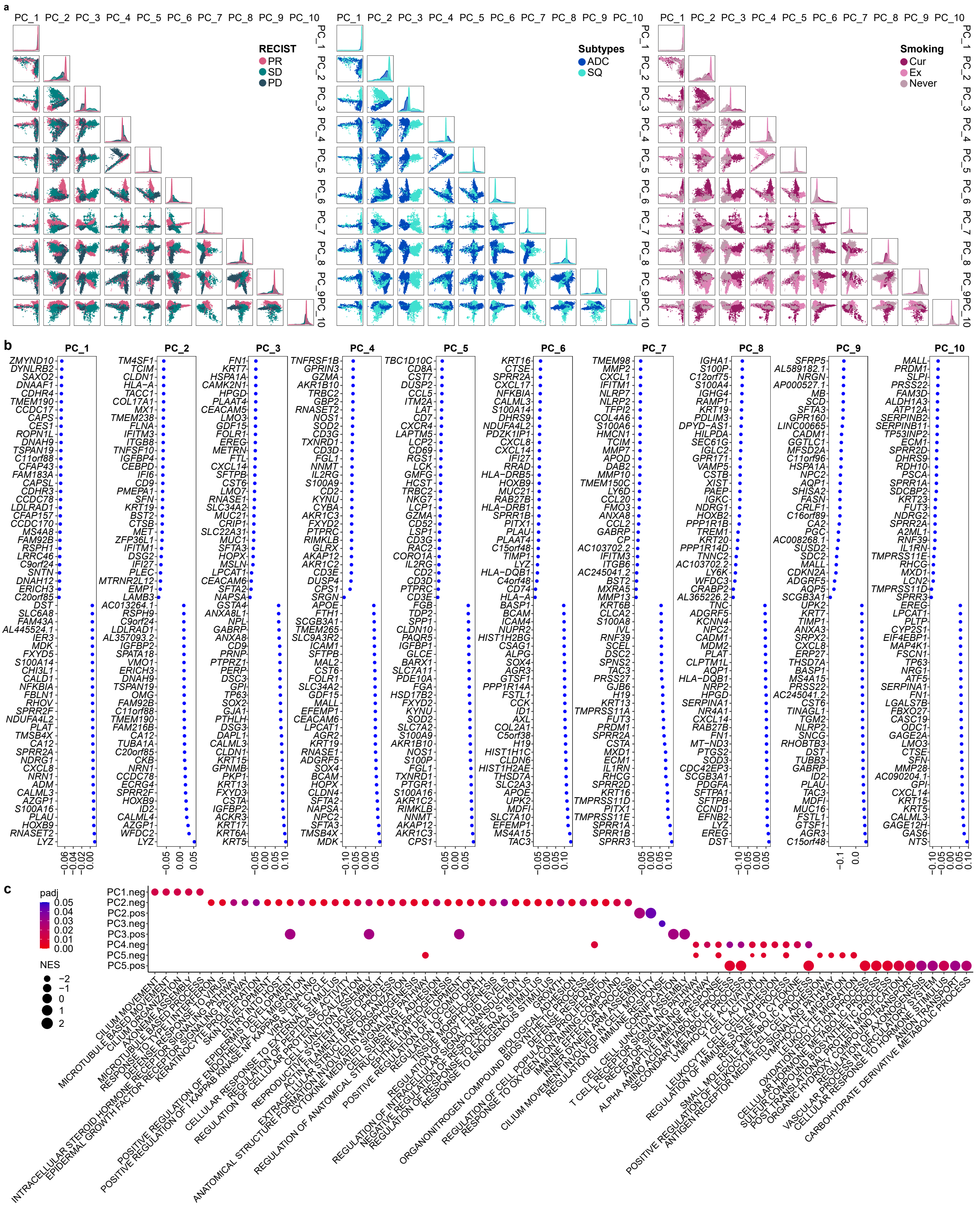


**Fig. S7. Selection and characteristics of PC signatures in malignant cells. a,** Unsupervised PCA projections of 12,975 malignant cells from PC1 to PC10, colored by RECIST, cancer subtypes, and smoking status. **b,** Dot plot of top 60 (30 genes with + and – scores) genes that contributed to each PC. **c,** Enrichment map of significant GO terms for each PC signature. Color and size indicate the adjusted p-value (padj) and the normalized enrichment score (NES) calculated using fgsea R package, respectively.

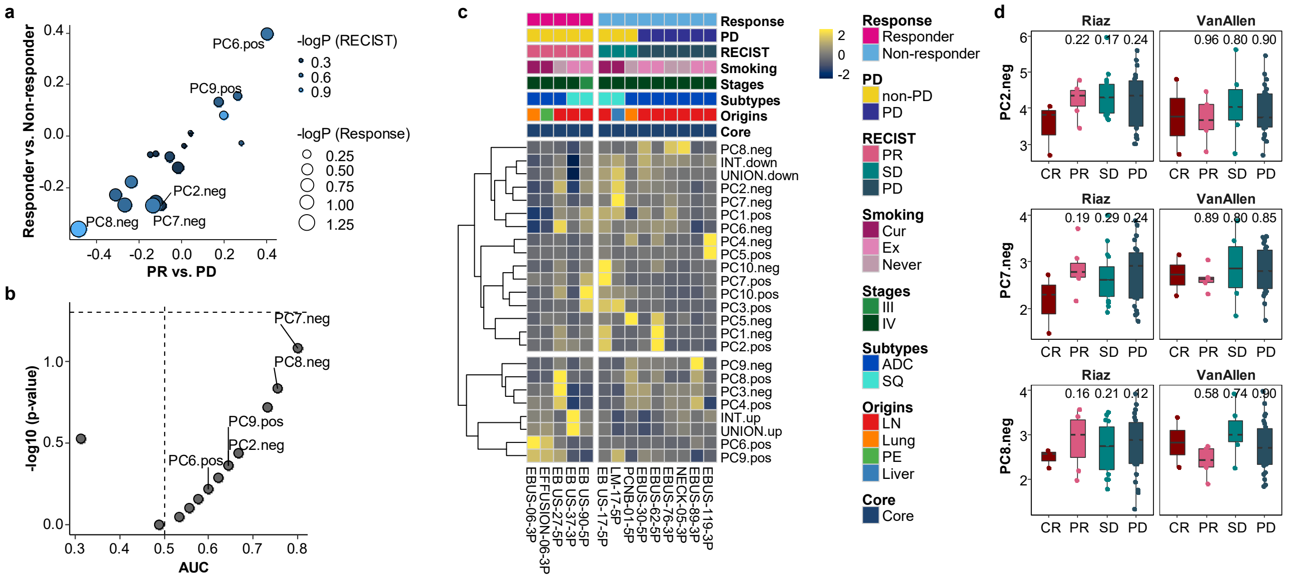

**Fig. S8. ICI response association of PC signatures in malignant cells. a,** Comparison of mean expression difference for each PC signature along ICI responses. Dot size and color represent -log (p-value) for responder vs. non-responder and PR vs. PD, respectively. p-value, two-tailed Student's t-test. **b,** ROC analysis of PC signatures to classify responder and non-responder. p-value, two-tailed Wilcoxon Rank Sum test. **c,** Heat map of relative expression of PC signatures across individual samples aligned with clinical data. Mean expression of each PC signature in each sample is scaled by z-transform and visualized in the range from -2.5 to 2.5. INT and UNION, intersection and union of DEGs for responder vs. non-responder, PR vs. PD, and PR vs. SD in Fig. 4a. **d,** Box plot of mean expression of PC2.neg, PC7.neg, and PC9.neg genes in Riaz et al.'s and Van Allen al.'s melanoma ICI cohorts. Label represents p-value for comparisons of each group to complete response (CR), calculated using two-tailed Student's t-test. Each box represents the median and the IQR, and whiskers indicate the 1.5 times of IQR.
